# Integrative molecular and clinical profiling of acral melanoma identifies LZTR1 as a key tumor promoter and therapeutic target

**DOI:** 10.1101/2021.04.20.440286

**Authors:** Farshad Farshidfar, Cong Peng, Chaya Levovitz, James Knight, Antonella Bacchiocchi, Juan Su, Kahn Rhrissorrakrai, Mingzhu Yin, Mario Sznol, Stephan Ariyan, James Clune, Kelly Olino, Laxmi Parida, Joerg Nikolaus, Meiling Zhang, Shuang Zhao, Yan Wang, Gang Huang, Miaojian Wan, Xianan Li, Jian Cao, Qin Yan, Xiang Chen, Aaron M. Newman, Ruth Halaban

**Affiliations:** Institute for Stem Cell Biology and Regenerative Medicine, Stanford University, Stanford, CA, USA; Department of Biomedical Data Science, Stanford University, Stanford, CA, USA; Xiangya Hospital, Central South University, Changsha, China; IBM Research, Yorktown Heights, NY, USA; Yale Center for Genome Analysis, Yale University, New Haven, CT, 06520, USA; Department of Dermatology, Yale University School of Medicine, New Haven, CT, USA; Department of Pathology, Yale University School of Medicine, New Haven, CT, USA; Department of Internal Medicine, Section of Medical Oncology, Yale University School of Medicine, New Haven, CT, USA; Department of Surgery, Yale University School of Medicine, New Haven, CT, USA; Department of Molecular and Cellular Physiology, Yale University School of Medicine, New Haven, CT, USA; Department of Dermatologic Surgery Institute of Dermatology, Chinese Academy of Medical Sciences & Peking Union Medical College, Nanjing, China; Department of Bone and Soft Tissue oncology, Hunan Cancer Hospital, Affiliated Tumor Hospital of Xiangya Medical School of Central South University, Changsha, Hunan, China; Department of Dermatology, The Third Affiliated Hospital, Sun Yat-sen University, Guangzhou, China; Tenaya Therapeutics, South San Francisco, CA, USA; Rutgers Cancer Institute of New Jersey and the Department of Medicine, Robert Wood Johnson Medical School, Rutgers University, New Brunswick, NJ, USA

**Author notes:** These authors contributed equally (Farshad Farshidfar, Cong Peng). Corresponding authors: Xiang Chen, Xiangya Hospital, Central South University, Changsha, China., Aaron M. Newman, Institute for Stem Cell Biology and Regenerative Medicine, Stanford University, Stanford, CA, USA., Ruth Halaban, Department of Dermatology, Yale University School of Medicine, New Haven, CT, USA.

## Abstract

Acral melanoma, the most common melanoma subtype among non-Caucasian individuals, is associated with poor prognosis. However, its key molecular drivers remain obscure. Here, we performed integrative genomic and clinical profiling of acral melanomas from a cohort of 104 patients treated in North America or China. We found that recurrent, late-arising amplifications of cytoband chr22q11.21 are a leading determinant of inferior survival, strongly associated with metastasis, and linked to downregulation of immunomodulatory genes associated with response to immune checkpoint blockade. Unexpectedly, LZTR1 – a known tumor suppressor in other cancers – is a key candidate oncogene in this cytoband. Silencing of LZTR1 in melanoma cell lines caused apoptotic cell death independent of major hotspot mutations or melanoma subtypes. Conversely, overexpression of LZTR1 in normal human melanocytes initiated processes associated with metastasis, including anchorage-independent growth, formation of spheroids, and increased levels of MAPK and SRC activities. Our results provide new insights into the etiology of acral melanoma and implicate LZTR1 as a key tumor promoter and therapeutic target.

## INTRODUCTION

Over the last two decades, a tremendous effort has been made to understand the genomic basis of melanoma. Collectively, these analyses have shown that sun-exposed melanomas harbor a large number of mutations and genomic rearrangements associated with ultraviolet (UV) radiation^1–6^. In contrast, acral melanomas, originating from hairless skin such as palms and soles, display a lower mutational burden, a higher rate of structural alteration, and poorer survival outcomes^2, 4, 7–17^. *BRAF* and *NRAS* are the most frequently affected oncogenes in acral melanomas but at a lower frequency compared to sun-exposed melanomas, whereas *KIT* mutations are more common in acral melanomas^14, 18–23^. Copy number variation (CNV) is a well-established feature of acral melanomas, contributing to aberrant regulation of several pathways affecting cell proliferation and gene expression. These include amplification of *CDK4*, *CCND1*, *MAPK1* and *NOTCH2;* loss of *CDKN2A* (p16^INK4^) and *NF1*; inactivation of *TP53*; modifications of chromatin regulators (e.g., *HDAC* amplification and loss of *ARID1A* and *ARID1B*); and alterations in *TERT*^11, 13, 14, 20, 23–25^. Despite these findings, attempts to treat acral melanoma with targeted inhibitors, such as CDK4/6 inhibitors, have failed^12^.

While most genomic studies of acral melanoma have been limited to relatively small clinical cohorts^1, 9, 13, 26, 27^, a recent whole genome analysis of 87 patients – 90% of which were of European ancestry – further confirmed the importance of structural rearrangements and copy number aberrations in this disease^15^. Given the predominance of acral melanoma in non-Caucasian populations^28, 29^ and the lack of effective targeted treatment options, large-scale genomic surveys of acral melanomas from ethnically diverse populations are needed.

Here, we applied whole exome (tumor/normal) and RNA sequencing to characterize acral melanomas from 104 patients treated in the United States (*n* = 37) and China (*n* = 67), most of whom had long-term follow-up data available. Through comparative genomic analysis with 157 sun-exposed melanomas, we identified novel molecular features of acral melanoma; generated the first prognostic map linking highly recurrent somatic aberrations in acral melanoma to risk of death; and found that late-arising focal amplifications in chr22q11.21 are associated with lymph node involvement and distant metastasis, leading to poor outcomes. Within chr22q11.21, we identified LZTR1 – a known tumor suppressor in other cancers – as a key candidate driver of metastasis. Our findings reveal novel molecular insights of acral melanoma pathogenesis and designate LZTR1 as a new therapeutic target.

## RESULTS

### Genomic characteristics of acral and sun-exposed melanoma

To characterize the genomic landscape of acral melanoma across ethnically diverse patient populations, we analyzed 104 tumors, including 97 by whole exome sequencing (WES), from patients treated in North America (‘Yale’) or China (‘CSU’) (Table 1). Both cohorts spanned all disease stages, included long-term follow-up, and encompassed patients with distinct ethnic origins, including Caucasian (*n* = 31; Yale) and Asian ancestry (*n* = 67; CSU) (**Supplementary Fig. 1a**). We also applied WES to profile 134 tumors from patients with sun-exposed melanoma, including patients with stage I through IV disease (Table 1). Notably, sun-exposed patients showed longer survival time than acral melanoma patients, consistent with previous studies^16, 17^ (**Supplementary Fig. 1b**). Peripheral blood leukocytes were analyzed as germline controls and whole-transcriptome sequencing (RNA-seq) was applied to 105 tumors, including 38 acral and 37 sun-exposed melanomas with matched WES data (Table 1). Tumor purities, clinical follow-up, and median survival times were comparable between acral cohorts, supporting their combined assessment (**Supplementary Fig. 1c,d**).

**Table 1:**
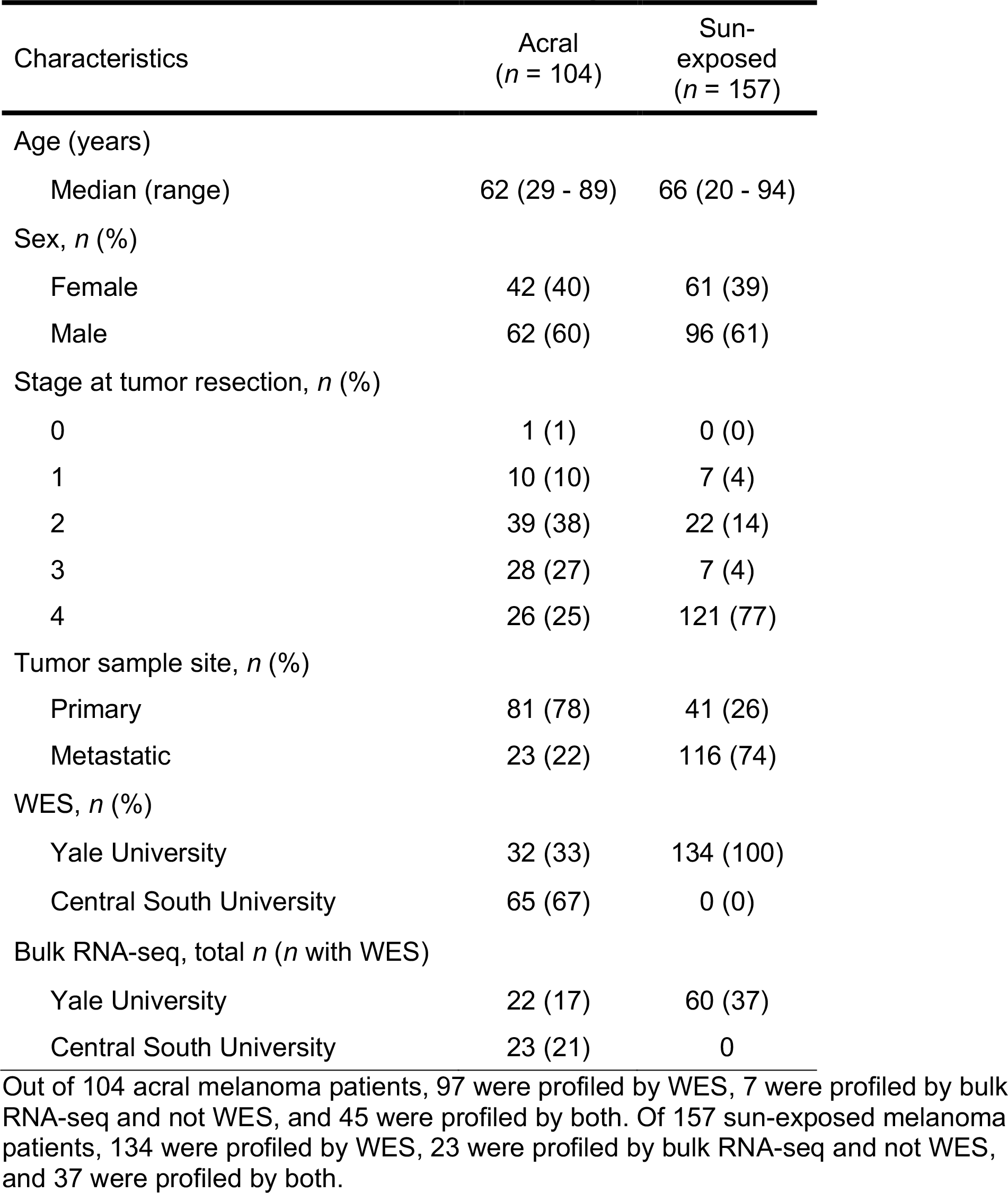
Patient characteristics and sequencing data.

| Characteristics | Acral<br>(*n* = 104) | Sun-exposed<br>(*n* = 157) |
| --- | --- | --- |
| Age (years) |  |  |
| Median (range) | 62 (29 - 89) | 66 (20 - 94) |
| Sex, *n* (%) |  |  |
| Female | 42 (40) | 61 (39) |
| Male | 62 (60) | 96 (61) |
| Stage at tumor resection, *n* (%) |  |  |
| 0 | 1 (1) | 0 (0) |
| 1 | 10 (10) | 7 (4) |
| 2 | 39 (38) | 22 (14) |
| 3 | 28 (27) | 7 (4) |
| 4 | 26 (25) | 121 (77) |
| Tumor sample site, *n* (%) |  |  |
| Primary | 81 (78) | 41 (26) |
| Metastatic | 23 (22) | 116 (74) |
| WES, *n* (%) |  |  |
| Yale University | 32 (33) | 134 (100) |
| Central South University | 65 (67) | 0 (0) |
| Bulk RNA-seq, total *n* (*n* with WES) |  |  |
| Yale University | 22 (17) | 60 (37) |
| Central South University | 23 (21) | 0 |

Out of 104 acral melanoma patients, 97 were profiled by WES, 7 were profiled by bulk RNA-seq and not WES, and 45 were profiled by both. Of 157 sun-exposed melanoma patients, 134 were profiled by WES, 23 were profiled by bulk RNA-seq and not WES, and 37 were profiled by both.

To verify key somatic lesions in acral melanoma, we began by performing a comparative genomics analysis. We observed striking variation in the prevalence of single nucleotide variants (SNVs) and insertions/deletions (indels) between melanoma subtypes, confirming a nearly ten-fold lower mutational burden in acral melanoma^2, 13^ (median of 406 vs. 42 nonsynonymous variants per exome in sun-exposed *vs*. acral melanoma, respectively; *P* = 2.2 x 10^-6^, two-sided Wilcoxon rank sum test; Fig. 1a). The most commonly mutated genes in acral melanomas were RAS family members (22% in *NRAS*, *KRAS*, and *HRAS*), followed by *KIT* (15%), *CGREF1* (10%), *BRAF* (8%), and *TP53* (4%). With the exception of *CGREF1*, which was limited to CSU patients, recurrence frequencies were similar between cohorts. Mutational signature analysis^17^ corroborated the prevalence of UV-induced mutagenesis in sun-exposed melanomas. In contrast, mutational signatures in acral melanomas were largely attributable to deamination of 5-methylcytosine (signature 1), which can arise from reactive oxygen species during melanin synthesis^30^, as well as alkylating agents (signature 11) and APOBEC activity (signature 13) (Fig. 1a**, bottom**).

**Figure 1:**
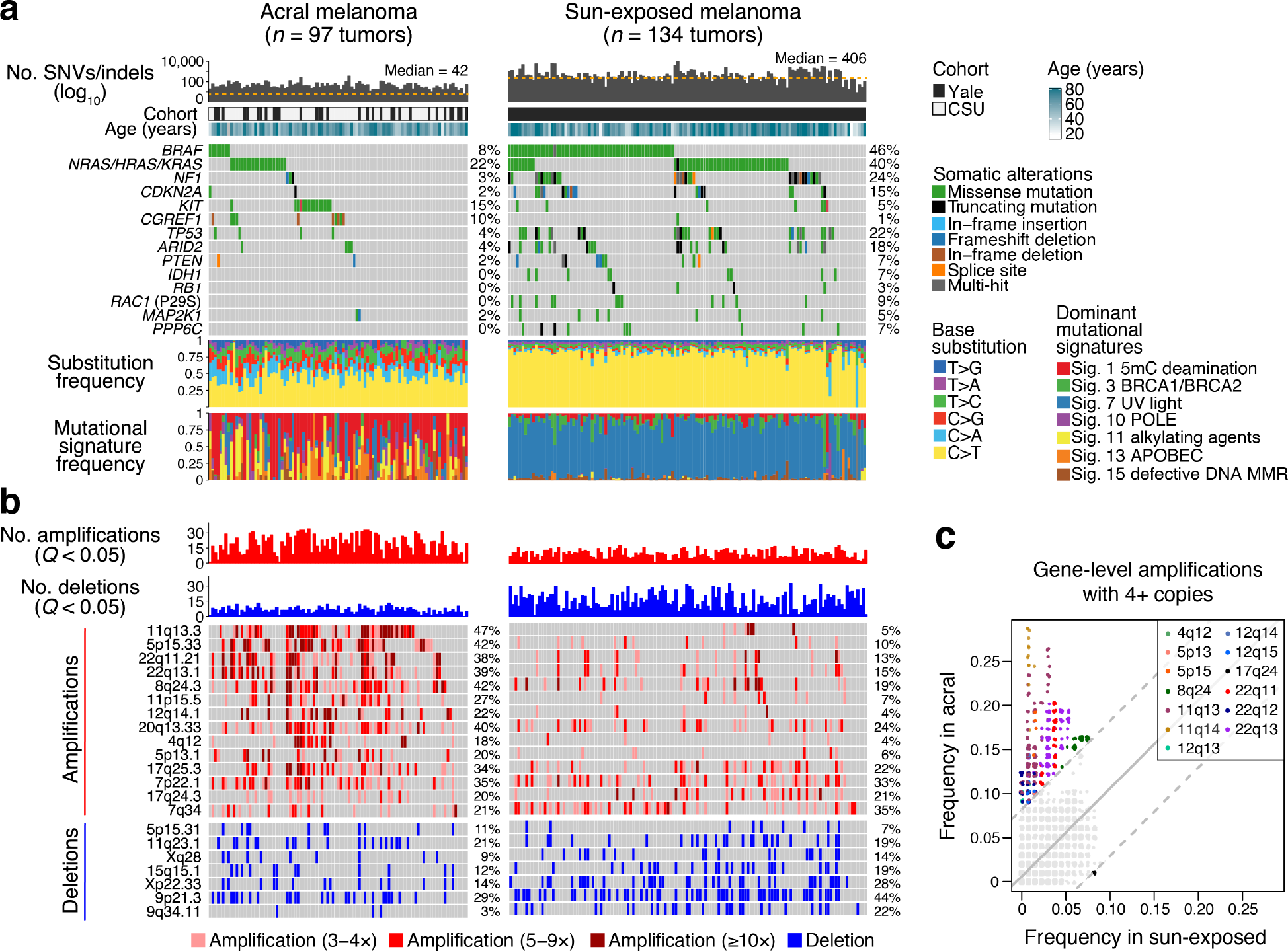
Landscape of somatic alterations in acral and sun-exposed melanomas. **a**-**b**, Genomic and clinical characterization of acral and sun-exposed melanoma samples sequenced in this work. **a,** The number of nonsynonymous SNVs and indels per melanoma exome (columns), cohort, age, frequently mutated genes (at least 5% recurrence frequency in either melanoma subtype), nonsynonymous base substitution frequencies, and dominant COSMIC mutational signatures^92, 93^. Sig., signature; 5mC, 5- methylcytosine. **b**, The number of significant focal amplifications and deletions (GISTIC *Q* < 0.05) per melanoma exome (columns), ordered identically to panel a. Cytobands with focal amplifications or deletions with at least 10% recurrence frequency in either melanoma subtype are shown (GISTIC *Q* < 10^−5^), ordered by the relative difference in recurrence frequency in acral versus sun-exposed melanoma. **c**, Genes are plotted according to the fraction of acral (*y*-axis) or sun-exposed (*x*-axis) tumors where they are present with 4+ copies. Considering the genome-wide distribution of differences in recurrence frequencies between melanoma subtypes, genes are identified as significantly recurrent in acral or sun-exposed melanomas if their |z-score| ≥ 3 (dashed lines). Significantly recurrent genes are colored according to their cytoband location (inset). For clarity, a small amount of jitter was added to distinguish overlapping genes.

As expected, focal amplifications were a core feature of acral melanoma in both cohorts (median of 20 *vs*. 11 per exome in acral *vs*. sun-exposed melanoma, respectively; *P =* 1.24 x 10^-10^, two-sided Wilcoxon rank sum test; **Fig.1b, top; Supplementary Figs. 1e** and **2a**). Among highly recurrent gene-level amplifications with at least four copies, those in chromosomes 4, 5, 8, 11, 12, and 22 were nearly exclusive to acral melanomas in our study (Fig. 1c). The most common amplifications enriched in acral melanoma were in cytobands 11q13.3 (47%), 5p15.33 (42%), 8q24.3 (42%), 22q13.1 (39%), and 22q11.21 (38%) (Fig. 1b; **Supplementary Fig. 2a**). Recurrent focal deletions, which included alterations in known genes such as *CDKN2A* (9p21.3)^22, 31, 32^, were less prevalent than in sun-exposed cases (**Fig.1b, bottom; Supplementary Fig. 2b**). Amplification and deletion frequencies were largely maintained in both acral cohorts.

Collectively, these results reveal somatic lesions in acral melanomas from genetically-distinct patient populations; corroborate and extend previous genomic studies^2, 4, 7–15^, and demonstrate the integrity and high quality of our data for downstream clinical analysis.

### Somatic determinants of risk in acral melanoma

Having systematically cataloged somatic aberrations in nearly 100 acral melanomas, we next sought to evaluate their clinical significance. We began by focusing on amplification events owing to their unique prevalence in this disease (Fig. 1b). Starting with the most statistically-significant peaks detected by GISTIC^33^ in a pooled analysis of both acral cohorts (*Q* < 10^−5^), we identified several loci linked to adverse overall survival, including peaks involving cytobands 22q11.21 and 22q13.1. Among them, cytoband 22q11.21 was most strongly associated with inferior overall survival (adjusted *P* < 0.05, univariate Cox regression of time from tumor resection; **Supplementary Fig. 3a**). This result was maintained when expanding the analysis to include all focal events identified by GISTIC (*Q* < 0.05) with at least 10% recurrence frequency in both acral cohorts and all genes with a nonsynonymous mutation frequency of at least 5% in either melanoma subtype (Fig. 2a). We also considered focal amplifications identified from the largest cohort (CSU) and tested in each cohort separately (Fig. 2b**; Supplementary Fig. 3b**). Again, 22q11.21 amplification was a leading determinant of adverse survival.

**Figure 2:**
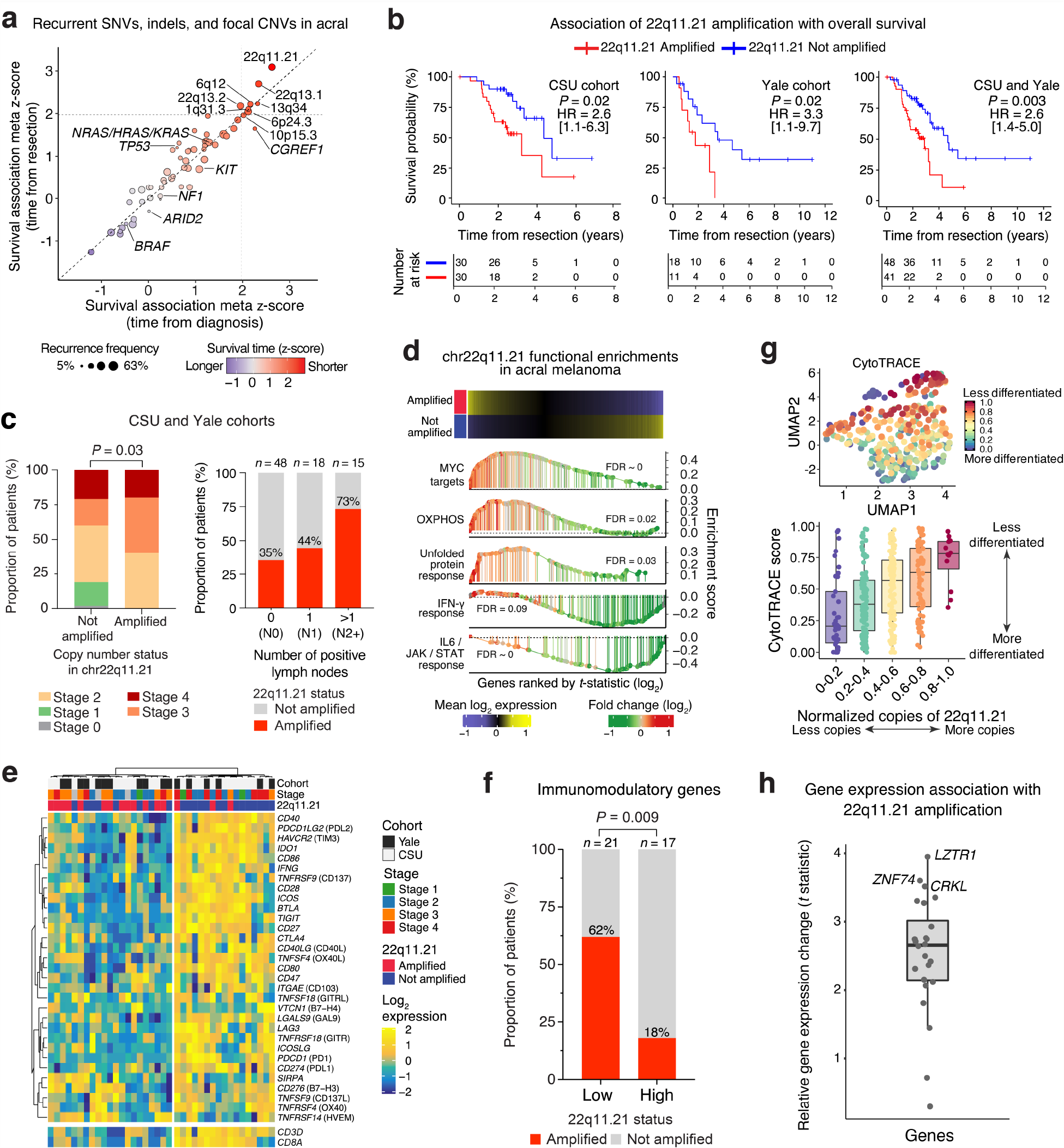
Focal amplifications in 22q11.21 are linked to shorter survival time, regional metastasis, and depletion of immunomodulatory programs in acral melanoma. **a**, Association between recurrent somatic alterations and overall survival (OS) in acral melanoma. Z-scores with positive and negative values indicated shorter and longer survival time, respectively (|*Z*| > 1.96 is significant at *P* < 0.05; Methods). OS was calculated from the date of diagnosis (*x*-axis) and the date of tumor resection (*y*-axis). Shown are genes with a nonsynonymous mutation frequency of at least 5% in either melanoma subtype and focal copy number events with at least 10% recurrence frequency in each acral cohort. Focal amplifications in cytobands significantly associated with OS and recurrently mutated genes are labeled. Additional details are provided in Methods. **b**, Kaplan Meier curves showing overall survival of acral melanoma patients, stratified by 22.q11.21 amplification status and calculated from the date of tumor resection. Significance was assessed with the log-rank test. HR, hazard ratio. 95% HR confidence intervals are shown in brackets. **c**, *Left*: Acral melanoma tumor stage shown as a function of 22q11.21 amplification status. Statistical significance was evaluated by a Chi *X*^2^ test*. Right:* Fraction of 22q11.21-amplified melanomas stratified by the number of involved lymph nodes (N stage). **d**, Hallmark signaling pathways significantly enriched in 22q11.21-amplified vs. non-amplified acral melanomas, as determined by pre-ranked Gene Set Enrichment Analysis (GSEA). Genes were ranked by log_2_ fold change adjusted for stage (Methods). OXPHOS, oxidative phosphorylation. **e**, Hierarchical clustering of 31 immunomodulatory genes (average linkage with Euclidean distance) in acral melanomas. *CD3D* and *CD8A* are included as lineage markers for T cells and CD8 T cells, respectively. **f**, Bar plot showing frequency of 22q11.21-amplified acral melanomas separated into high and low immunomodulatory expression groups, as defined by clustering in panel e. Statistical significance was determined by Fisher’s exact test. **g,** Single-cell differentiation status, as imputed by CytoTRACE^42^ (top), versus the estimated relative number of 22q11.21 copies per cell (normalized between 0 and 1), as imputed by CONICSmat^36^ (bottom), in cancer cells from a 22q11.21-amplified acral melanoma tumor (**Supplementary Fig. 4**; Methods). Single-cell transcriptomes are visualized in the top subpanel by Uniform Manifold Approximation and Projection (UMAP). **h**, Association between gene expression and 22q11.21 amplification status in Yale and CSU acral melanoma cohorts. Only genes physically located on cytoband 22q11.21 are shown. Group comparisons were performed using a two-sided *t*-test with unequal variance. The top three genes are indicated. Amplified, 3+ copies.

Given this observation, we sought to better understand 22q11.21 focal amplification and the factors underlying its clinical phenotype. We first tested whether 22q11.21 is a surrogate for advanced disease at the time of tumor resection. Intriguingly, 22q11.21 amplifications were observed across all stages except stage I disease (*P* = 0.03, Chi *X*^2^ test; Fig. 2c**, left**). We verified this result in three independent acral melanoma cohorts, including an external dataset comprised of 33 patients for whom stage at presentation was known^11^, demonstrating that 22q11.21 amplification is a recurrent late-arising event in acral melanoma (**Supplementary Fig. 3c**). Given this result, we reassessed survival associations using stage as a covariate. Regardless of whether we examined all patients or just those with stage II through IV disease, 22q11.21 amplifications remained significant after multivariate adjustment for stage (*P* = 0.008 and 0.024, respectively; Cox proportional hazards regression). This was also true for acral patients with advanced disease (III or IV) (*P* = 0.03, Cox proportional hazards regression; **Supplementary Fig. 3c**), for whom stage alone did not significantly stratify outcomes.

As a common late-arising event, we next tested if 22q11.21 amplifications might correlate with tumor progression. Indeed, in both acral cohorts, we observed a strong positive correlation between 22q11.21 amplification frequency and the number of positive lymph nodes (Fig. 2c**, right; Supplementary Fig. 3e**). Remarkably, nearly 75% of patients with >1 positive lymph node harbored at least one gain of 22q11.21 (Fig. 2c**, right**). Reanalysis of WES data from an independent study^11^ confirmed this trend (**Supplementary Fig. 3f**). While this association was observed in both primary and metastatic tumor specimens, the latter showed a modest but consistent increase in amplification frequency after controlling for lymph node status (**Supplementary Fig. 3g**). No other associations with clinical indices were observed (**Supplementary Fig. 3h**).

Taken together, these data reveal that 22q11.21 focal amplification is a conserved, late-arising somatic event linked to poor survival and regional metastasis in acral melanoma, independent of Caucasian or Asian ancestry. Accordingly, this event could represent a critical step in the initiation or maintenance of nodal metastasis.

### Integrative genomics of 22q11.21 focal amplification

To understand the biological significance of 22q11.21 amplification in acral melanoma, we next examined transcriptional hallmarks of 22q11.21-amplified tumors. By employing a linear model adjusted for stage (**Methods**), we rank-ordered genes by their differential expression in 22q11.21-amplified tumors and performed gene set enrichment analysis^34^ (Fig. 2d). Overall, 22q11.21-amplified melanomas were significantly enriched in canonical signaling pathways associated with tumorigenesis and metabolic activity, including MYC target genes, oxidative phosphorylation, and unfolded protein response^35^. In contrast, patients with non-amplified tumors showed higher expression of immunoreactive programs such as IL6/JAK/STAT and IFN-γ response pathways. We hypothesized that such patients might be superior candidates for existing or emerging immunotherapies (Fig. 2d). Consistent with this possibility, we observed a striking reciprocal relationship between 22q11.21-amplification and the expression of immunomodulatory genes, including key targets of immune checkpoint blockade (e.g., *PDCD1*, *CTLA4*) (Fig. 2e). Among patients with high expression of immunomodulatory genes, only 12% were amplified, whereas among patients with low expression, 62% were amplified (Fig. 2f). This result was highly significant (*P* = 0.009, Fisher’s exact test), indicating that 22q11.21-amplified and non-amplified tumors preferentially reflect “cold” and “hot” tumor microenvironments, respectively.

To extend these observations to single cells, we applied single-cell RNA sequencing (scRNA-seq) to an acral melanoma tumor specimen with four additional copies of 22q11.21, as determined by WES (**Supplementary Fig. 4a**). Using canonical marker genes and copy number inference via CONICSmat^36^, 321 single-cell transcriptomes were confidently identified as melanoma cells (**Supplementary Fig. 4b, Methods**). We confirmed over-expression of genes on the 22q arm, consistent with WES (**Supplementary Fig. 4c**). However, 22q expression levels were heterogeneous across cells, indicating variability in the number of copies per cell (**Supplementary Fig. 4c**). By dividing malignant cells into two groups according to the median expression of 22q arm genes (selected from within the 22q11.21 peak identified by GISTIC), we observed the same amplification-enriched pathways identified in bulk tumors, including oxidative phosphorylation and MYC targets, confirming their malignant origin (**Supplementary Fig. 4d**).

Immature cancer cells often display elevated metabolism via oxidative phosphorylation and MYC activity^37^ and stemness features in melanoma tumors have been linked to poor survival^38–41^. To test whether 22q11.21-amplified cells exhibit an immature cellular phenotype, we employed CytoTRACE, a recently described *in silico* method for predicting developmental potential on the basis of single-cell transcriptional diversity^42^. Indeed, cells with higher relative copies of 22q11.21 were predicted to be less mature (Fig. 2g). Notably, this result was independent of genes physically located on 22q, implying that 22q11.21-amplified cells exhibit a more accessible genome, a hallmark of immature cells in normal tissues^42^ (**Supplementary Fig. 4e**).

Finally, we used *t-*statistic to rank genes in 22q11.21 according to their expression in amplified versus non-amplified tumors (Fig. 2h). The top-ranking gene associated with amplification was *LZTR1* (leucine zipper like transcription regulator 1). We were struck by this result because LZTR1, a member of the Kelch-like (KLHL) family and an adaptor for Cullin 3 (CUL3) ubiquitin ligase complexes^43, 44^, is considered a tumor suppressor in schwannoma and glioblastoma^43, 45, 46^. Nevertheless, we found that high expression of *LZTR1* is predictive of poor outcome, both in acral and sun-exposed melanomas from this study, and in 443 advanced sun-exposed melanomas profiled by TCGA (The Cancer Genome Atlas) (**Supplementary Fig. 5, Methods**). Beyond *LZTR1*, we noted that *ZNF74*, a zinc finger protein, and *CRKL* (CRK like proto-oncogene, adaptor protein), a recurrently amplified gene in multiple carcinomas^47–50^, including non-small cell lung cancer (3% – 13% of cases)^47–49^, were ranked 2^nd^ and 3^rd^ in our analysis, respectively. Given these results, we set out to characterize the biological functions of these genes to determine which, if any, underlie the observed clinical phenotype of 22q11.21 amplification.

### Suppression of LZTR1 attenuates melanoma cell proliferation and induces apoptosis independent of Ras or MAPK activity

We began by silencing several chr22q11.21-amplified genes using lentiviral delivery of short hairpin RNAs (shRNAs), with the goal of determining the impact of targeted knockdowns on melanoma cell proliferation. Treatment of two acral melanoma cell lines with *ZNF74* shRNA had a modest effect on cell proliferation (**Supplementary Fig. 6a**). Similarly, while downregulation of *CRKL* induced growth arrest, only one of three shRNAs against *CRKL* successfully downregulated *CRKL*, and only two of five tested cell lines were highly affected (**Supplementary Fig. 6b**). Conversely, silencing of *LZTR1* consistently arrested cell proliferation. This was the case regardless of subtype (acral or sun-exposed) or mutations in *BRAF* or *NRAS* (Fig. 3a, b). In addition, we observed growth arrest in normal melanocytes derived from two independent foreskins (Fig. 3a, b). We ruled out off-target effects because six different LZTR1-directed shRNAs induced growth arrest, as did CRISPR-Cas9 sgRNA directed against LZTR1 (Fig. 3a-c; **Supplementary** Fig 5c). The observed phenotype had a long-term effect since LZTR1*-*null melanoma cells did not survive *in vitro*, whereas cells infected with control shRNA (scrambled) continued to proliferate. We also tested depletion of *SNAP29* and *THAP7*, both of which are physically located on 22q11.21 but whose expression levels were not significantly linked to 22q11.21 amplification. Knockdown of these genes had little to no effect on proliferation (**Supplementary Fig. 5d, e**).

**Figure 3:**
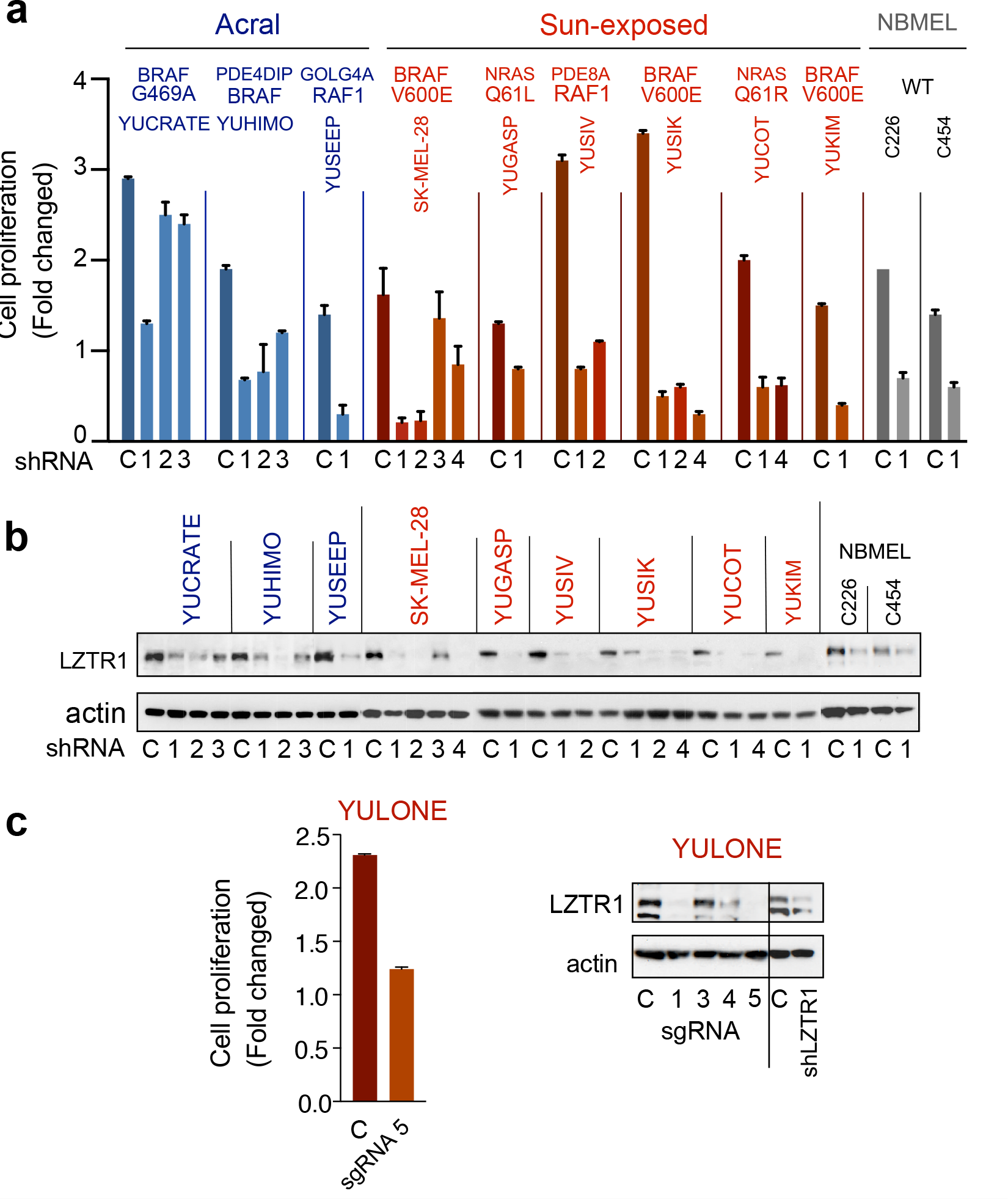
Cell proliferation in response to suppression of LZTR1. **a**, Impact of LZTR1 knockdown on cell proliferation in nine primary melanoma cell lines and two normal human melanocyte lines (NBMEL). Key mutations are indicated. Bar plots depict fold change between the 3^rd^ and 6^th^ day after infection with LZTR1 shRNA (numbered), as compared to control (scrambled) shRNA (‘C’). All LZTR1 shRNAs significantly reduced proliferation relative to control (*P* < 0.05; two-sided *t* test with unequal variance). **b**, Western blot showing the efficiency of LZTR1 knockdown in primary acral and sun-exposed melanoma cell lines, and in normal melanocytes, related to panel a. **c**, Cell proliferation (left) and LZTR1 expression (right) of a sun-exposed melanoma cell line that lost one LZTR1 allele in response to genomic modification by different CRISPR-Cas9 sgRNAs targeting LZTR1. Bar plots depict fold change between the 3^rd^ and 6^th^ day after infection with sgRNA 5. Reduced cell proliferation is statistically significant (*P* <0.05; two-sided *t* test with unequal variance). Bars in a, c represent the mean of triplicate or quadruplet wells and error bars indicate SEM. Actin levels in b and c show protein loading. Cell lines are indicated above all plots in a–c and colored according to their origin: acral melanoma (blue), sun-exposed melanoma (red), normal melanocyte (grey). NBMEL, newborn melanocyte cultured from Caucasian foreskin; WT, wildtype.

Given these results, we sought to better understand the biological consequences of LZTR1 knockdown. Inactivating germline mutations in LZTR1 are associated with Noonan syndrome and functional studies have linked LZTR1 inactivation to RAS ubiquitination, increased RAS-MAPK signaling, and cell proliferation^51–56^. Indeed, suppression of LZTR1 in melanoma cells increased the constitutive levels of GTP-bound RAS, an effect similar to that observed in growth factor-stimulated cells^54, 57^. RAS-GTP levels increased in NRAS- or BRAF-mutant melanoma cells without a change in total RAS protein (Fig. 4a, b).

**Figure 4:**
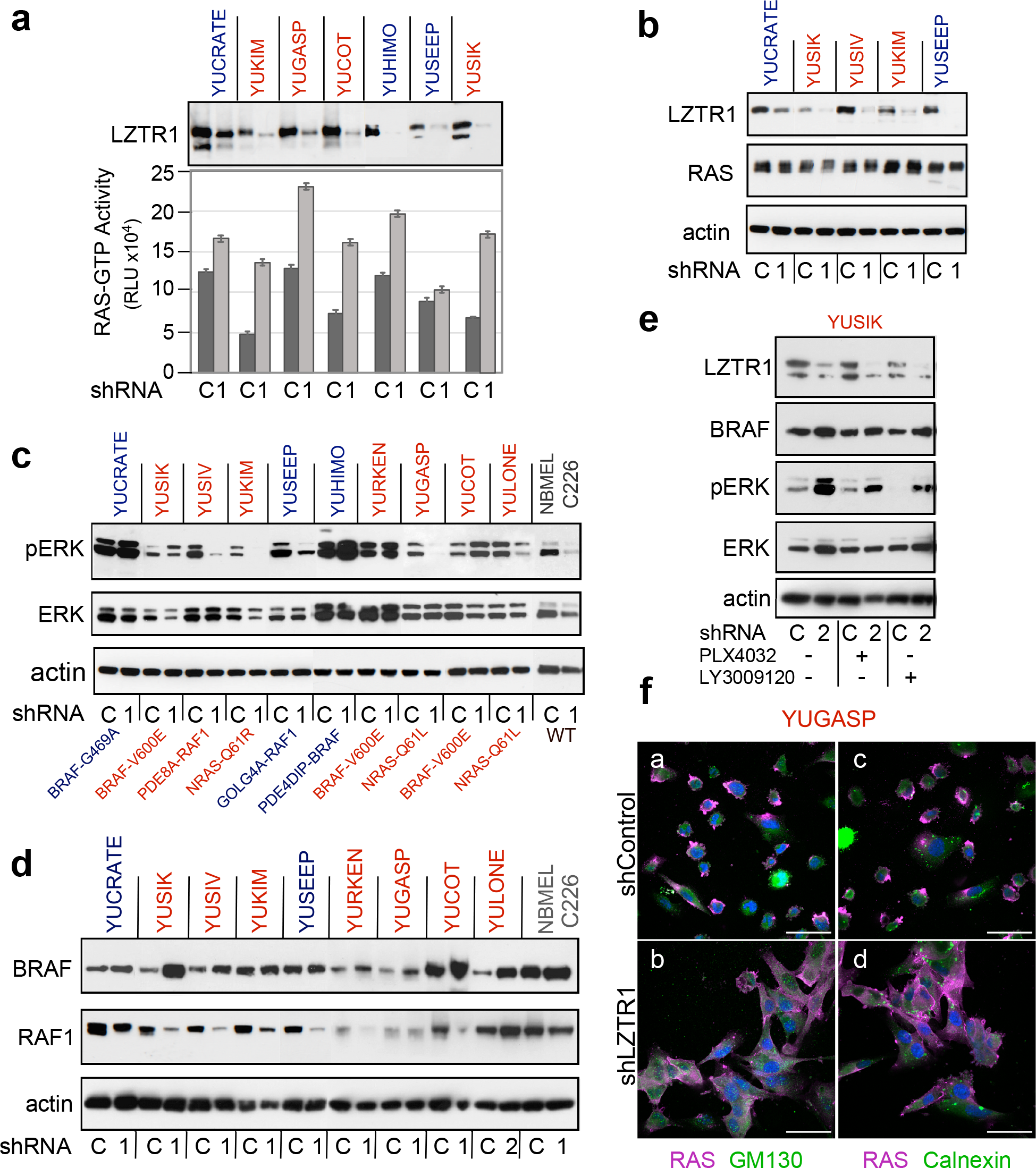
Changes in MAPK signaling in response to LZTR1 knockdown. **a, b**, Impact of LZTR1 loss on (**a**) RAS-GTP activity as measured by RAS-GTPase Activation ELISA assay, and (**b**) RAS levels, five days after shLZTR1 infection. The data in a represent average of triplicate or duplicate wells ± SE. **c**, Effect of LZTR1 loss on MAPK activity. Key somatic events are indicated below. WT, wildtype. **d, e** Increased levels of BRAF activity in response to shLZTR1 is associated with increase pERK. In panel e, cells were incubated with RAF the kinase inhibitors PLX4032 (500 nM) or LY3009120 (100 nM) for four hours at the end of treatment with shRNA. **f**, RAS translocation to the cytoplasm in response to shLZTR1; a-d, RAS is visualized by staining with magenta; Green (Cy2) indicates GM130 (a, b) and calnexin (c, d). Scale bar = 50 µm. Blue and red indicate acral and sun-exposed melanoma cell lines, respectively. shRNAs in a–e are indicated by numeric identifiers. C, scrambled shRNA control. Actin levels in a–e show protein loading. Cell lines are indicated above all plots in a–f and colored according to their origin: acral melanoma (blue), sun-exposed melanoma (red), normal melanocyte (grey). NBMEL, newborn melanocyte.

We also observed widespread changes in MAPK signaling following LZTR1 knockdown. For example, there was an increase in pERK in melanoma cells carrying *BRAF^V^*^600E^ or *PDE4DIP-BRAF* (Fig. 4c). In contrast, pERK decreased in *NRAS^Q61L/R^* melanoma cells, *GOLG4A*-*RAF1* or *PDE8A*-*RAF1* fusion-bearing melanoma cells, and normal human melanocytes (Fig. 4c). ERK activation was likely due to an increase in BRAF levels (Fig. 4d), enhancing BRAF activity. Treatment of melanoma cells with BRAF^V600E/K^ or pan-RAF inhibitors (PLX4032 or LY3009120) reduced shLZTR1-induced pERK activation (Fig. 4e), rendering further support for the role of BRAF kinase activity. On the other hand, ERK inhibition in shLZTR1-treated cells could potentially arise from RAS translocation to the cytoplasm (Fig. 4f), and the consequent disassociation from its membrane-bound mitogenic effectors, which are critical for *NRAS^Q^*^61^*^/L/R^* mutant and WT cells lacking BRAF mutations. RAS translocation was not linked to de-ubiquitination, because loss of LZTR1 did not change the levels of ubiquitinated RAS (**Supplementary Fig. 7a**). RAF1 levels were diminished in most cell lines (Fig. 4d), reflecting a decrease in gene expression. Thus, downregulation of LZTR1 induces growth arrest independently of ERK activity, the presence of BRAF or NRAS oncogenes, and changes in RAS-GTP levels.

We next explored if our *in vitro* melanoma systems effectively recapitulate key 22q11.21-related signaling pathways observed *in vivo*. To this end, we performed bulk RNA-sequencing of a melanoma cell line (YUSIK) to assess the impact of LZTR1 knockdown. Remarkably, depletion of LZTR1 induced transcriptome-wide changes that largely mirrored those observed in bulk tumors and single melanoma cells (Fig. 5a).

**Figure 5:**
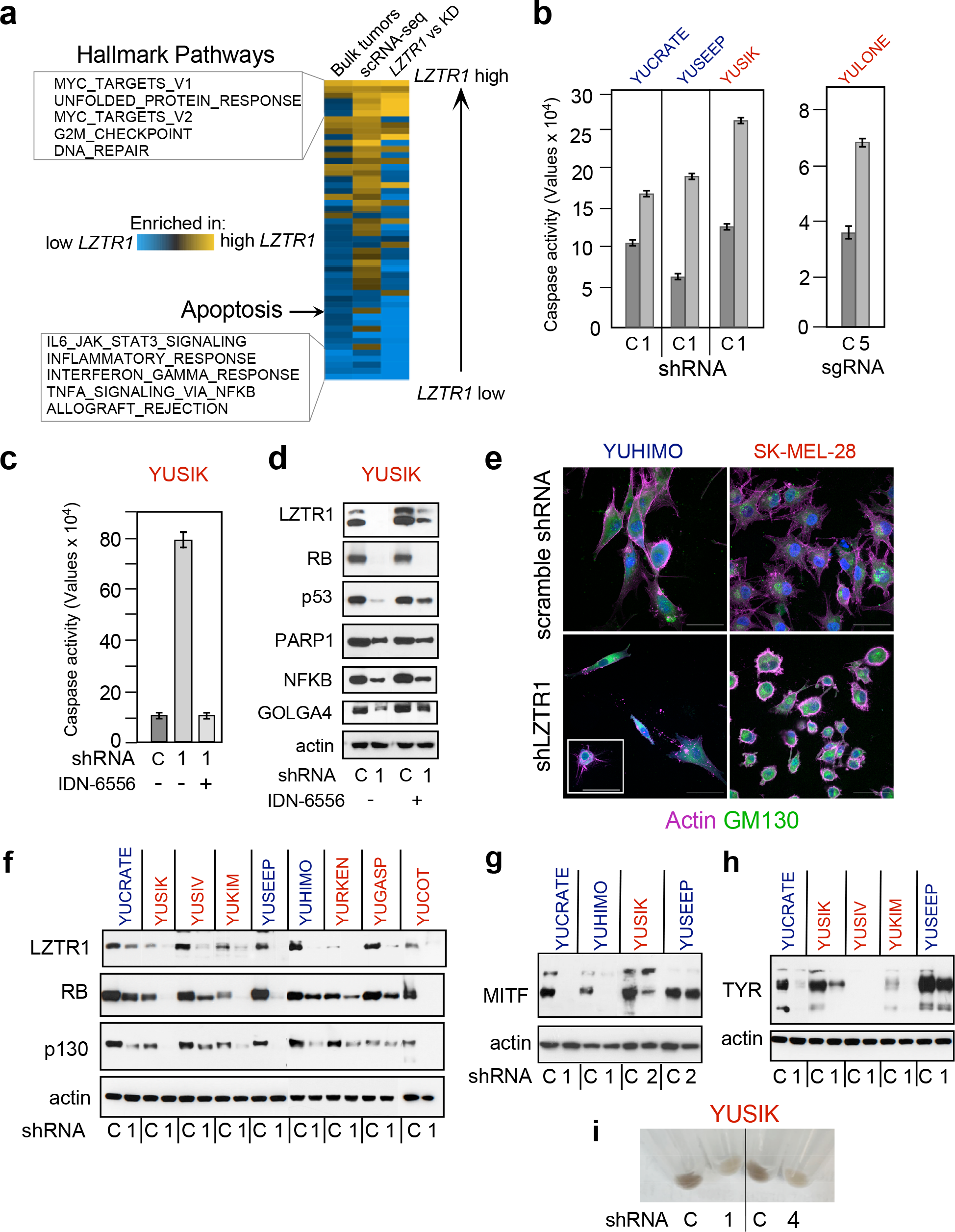
Impact of LZTR1 knockdown on apoptosis, Rb signaling, and pigmentation. **a**, Gene Set Enrichment Analysis (GSEA)^94^ showing concordance in hallmark pathways among bulk acral melanoma tumors, acral melanoma single-cell transcriptomes (scRNA-seq), and a primary melanoma cell line (*LZTR1* vs. KD), in relation to high vs. low *LZTR1* expression. Gold, high positive normalized enrichment score (NES); blue, high negative NES; KD, knockdown. **b,** LZTR1 shRNA and sgRNA (CRISPR-Cas9) induce apoptosis in melanoma cells. **c, d,** Effects of inhibiting caspase activity with IDN-6556 (IDN, 2 µM, 3 days). As shown in c, IDN-6556 suppressed shLZTR1-induced caspase activity. As shown in d, IDN-6556 increased the levels of LZTR1 (known to be degraded by caspases) and rescued caspase substrates, such as GOLGA4, p53, and to a lesser extent, NF-κB. **e**, Effect of shLZTR1 on cell morphology and actin filament organization. Actin filaments were visualized by staining with rhodamine-phalloidin (magenta) and the Golgi with anti-GM130 (green, Cy2). The nuclei are stained with DAPI (blue). Scale bar = 50 µm. **f**, shLZTR1 downregulates RB and p130 (also known as RBL2). **g–i,** Impact of shLZTR1 on MITF (panel g), tyrosinase (TYR) (panel h), and pigmentation (panel i). shRNAs in b-d and f-i are indicated by unique numerical identifiers. C, scrambled shRNA control. Actin levels in **d** and **f-h** show protein loading. Of note, the actin control in f and h is identical because the same membrane was used to blot the proteins. Cell lines are indicated above all plots in **b–g** and colored according to their origin: acral melanoma (blue), sun-exposed melanoma (red).

We noticed that among altered transcriptional programs, apoptosis-related genes were elevated in cell lines and tumors with lower LZTR1 expression (Fig. 5a). These data are supported by an increase in caspase activity after treatment with LZTR1 shRNA or sgRNA (Fig. 5b, c), which led to the degradation of known caspase substrates^58^, including pRb, p53, PARP1, NFKB, and GOLGA4 (Fig. 5d). Notably, GOLGA4 localizes to the Golgi apparatus, the subcellular site of LZTR1^59^. Moreover, shLZTR1-induced caspase activity was suppressed by the pan-caspase inhibitor IDN-6556 (Emricasan), which also rescued several substrates, including LZTR1 (Fig. 5c, d). These data are consistent with a previous report showing that LZTR1 undergoes caspase-mediated degradation^59^. Furthermore, shLZTR1 led to disruptions of cellular organization, including actin depolymerization into irregular shapes (Fig. 5e, left), or formation of actin rings around the Golgi and nucleus (Fig. 5e, right). Such changes are characteristic of cells undergoing fast or slow apoptotic death, respectively^60^.

Several cell cycle proteins were also downregulated, in line with pathway enrichment analyses (Fig. 5f and **Supplementary Fig. 7b-e**). In addition, retinoblastoma proteins (pRb and p130) were suppressed in the nine melanoma cell lines tested (Fig. 5f). This was likely due to ubiquitination and degradation^61^, as pRb was not rescued by the caspase inhibitor IDN-6556 (Fig. 5d). Elimination of pRb may enhance mitochondrial-mediated apoptosis because it leads to reduced mitochondrial mass, reduced activity of the electron transport chain, and increased reactive oxygen species (ROS)^62–65^.

A major reason for growth arrest in some melanoma cell lines is downregulation of MITF, a lineage-specific transcription factor critical for melanocyte and melanoma cell proliferation^66^. MITF stability is reduced when phosphorylated by MAPK or KIT^67, 68^, and this process was clearly observed in three out of four melanoma cell lines with increased ERK activity (Fig. 5g, as compared to Fig. 4c, e). Downregulation of MITF, as expected, is associated with decrease in tyrosinase (TYR), the key enzyme in melanin synthesis as well as cellular pigmentation (Fig. 5h, i). These results are consistent with our published observations using the same melanoma cell lines^69^.

### Overexpression of LZTR1 in normal melanocytes confers properties of malignant transformation and metastasis

We next evaluated the impact of overexpressing LZTR1 in normal melanocytes and compared the effects to overexpression of CRKL. The latter is a SH3/SH2 adaptor protein that promotes lung cancer cell invasion via ERK activation^70^ and epithelial– mesenchymal transition (EMT) in colorectal and pancreatic carcinomas^71^. Early passage human melanocytes (passage 4) were transduced with HA-tagged *LZTR1* cloned into the pInd20 lentiviral vector, V5-tagged *CRKL* inserted into the PLX304 vector, or both constructs. Over-expression of these genes did not enhance the rate of cell proliferation; rather, melanocytes overexpressing CRKL grew slower compared to parental cells (**Supplementary Fig. 8a**). Nevertheless, within 2-3 days after infection, we noticed a striking induction of anchorage-independent growth, observed as cells overexpressing LZTR1 or CRKL formed three-dimensional clusters in 2D and 3D collagen cultures (Fig. 6a, top and bottom rows, respectively). Moreover, this result – which was reminiscent of a malignant cell phenotype^72^ – was further enhanced when both genes were co-expressed (LZTR1+CRKL), leading to the formation of spheroids that detached from the surface of the dish (Fig. 6a).

**Figure 6:**
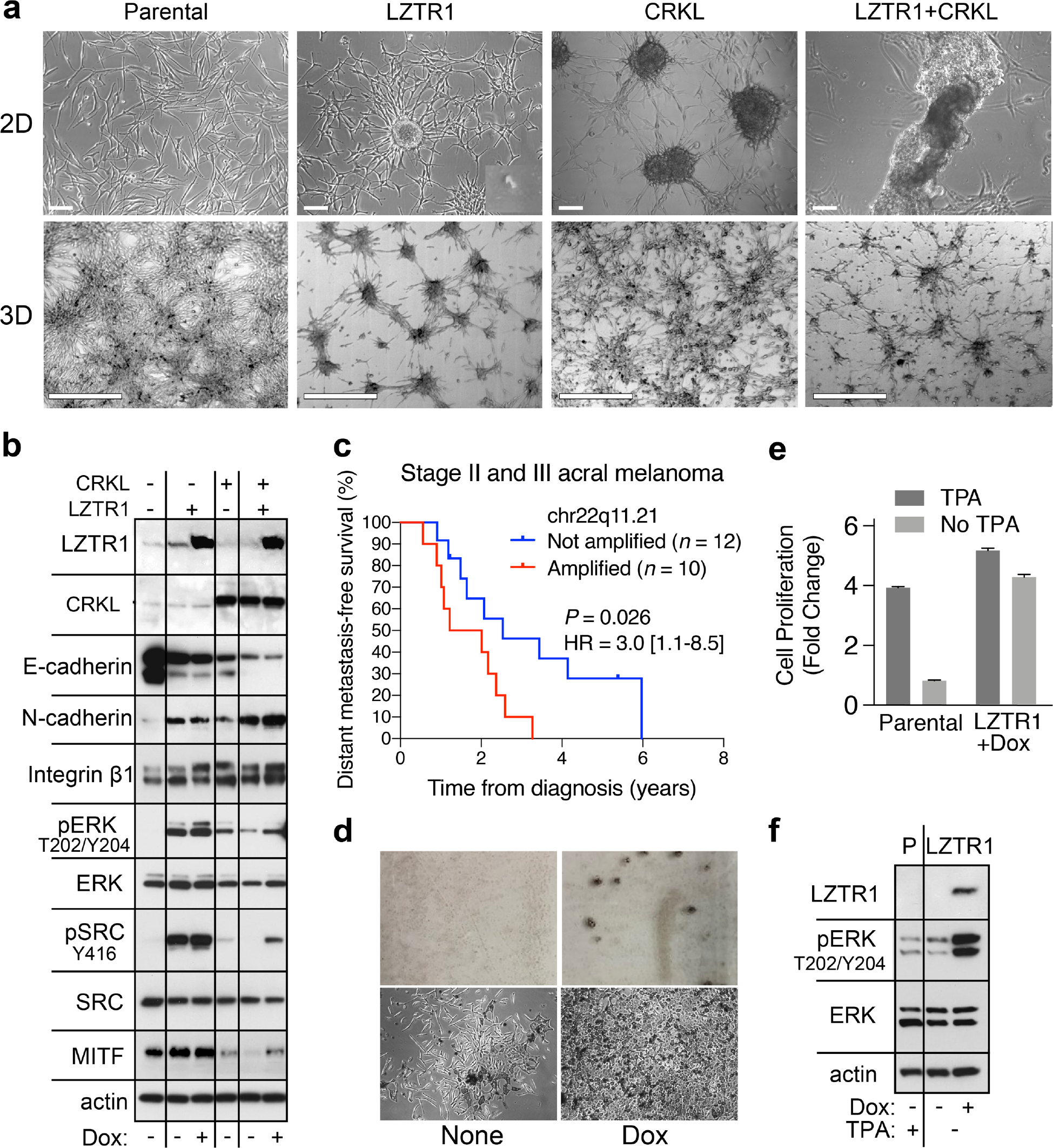
LZTR1 and CRKL confer properties consistent with malignant cell transformation and metastasis initiation. **a**, Morphological changes and spheroid formations in early passage normal human melanocytes (NBMEL C1220) overexpressing LZTR1 and/or CRKL. *Top*: Phase-contrast images of parental and infected cells in 2D culture. LZTR1 images were taken after two days induction with doxycycline (200 ng/ml), CRKL after three days of infection with PLX304-CRKL, and LZTR1+CRKL after six days infection of LZTR1 melanocytes with PLX304-CRKL and three days stimulation with doxycycline. Insert in LZTR1 shows that colonies were evident without magnification. The non-induced LZTR1 cells grow as a monolayer, as seen for the parental non-transformed melanocytes (Parental). Scale bar = 100 µm. *Bottom*: Low magnification images showing 3D cultures of melanocytes seeded in 0.5% collagen for three days. LZTR1 and CRKL, both alone and in combination, induced aggregation and multicellular spheroids. Scale bar = 500 µm. **b**, Western blot showing critical changes in normal human melanocytes overexpressing LZTR1, CRKL, or both (as seen in the top two lanes) compared to parental (–). Cells were harvested after incubation in regular medium, or medium supplemented with doxycycline for two days when applicable (Dox, 200 ng/ml). Of note, an increase in LZTR1 produced by basal promoter activity was sufficient to induce constitutive MAPK and SRC activities. Actin levels show protein loading. **c**, Kaplan Meier plot showing differences in distant metastasis-free survival (DMFS) between acral melanoma patients stratified by 22q11.21-amplification status. Patients with stage II or III disease at diagnosis with available DMFS data are shown (Yale cohort). Statistical significance was assessed by a log-rank test. HR, hazard ratio. 95% HR confidence interval is shown in brackets. **d**, 2D cultures of spontaneously immortalized mouse melanocytes (C57BL) forming colonies in the absence of TPA in response to LZTR1 (Dox). The dark colonies are seen without magnification (top), and under phase-contrast microscope (bottom). **e**, Cell proliferation of parental and LZTR1 transformed C57BL mouse melanocytes. **f**, Western blots displaying LZTR1 expression and MAPK activation (pERK) in TPA-starved (–) mouse melanocytes in response to doxycycline (+, 200 ng/ml), compared to parental, non-transformed cells (P).

During metastasis, primary melanoma cells detach from the dermis and migrate to secondary sites through increased cell-cell interactions and promotion of cancer cell survival. We therefore examined changes in adhesion proteins affecting cell-matrix and cell-cell interactions known to mediate aggregation, the formation of spheroids^72^, and *in vivo* EMT^73, 74^. Our data show that E-cadherin was downregulated whereas N-cadherin and integrin β1 were upregulated in response to increased expression of LZTR1 and CRKL, a process that was enhanced when both genes were co-expressed (Fig. 6b). Notably, our results with CRKL were consistent with HCT116 colon cancer cells, in which loss of CRKL was found to increase E-cadherin expression and shift the cells toward an epithelial phenotype^71^. Importantly, LZTR1 and CRKL, both alone and in combination, induced high levels of constitutively active ERK and SRC relative to parental cells (Fig. 6b, pERK and pSRC), functions that support viability and proliferation. We also identified downregulation of MITF in cells overexpressing CRKL as the possible cause for growth rate attenuation (Fig. 6b; **Supplementary Fig. 8a**). Consistent with this finding, while higher expression of MITF defines a proliferative subtype of melanoma (MITF^high^–AXL^low^), lower expression is preferentially associated with invasion (MITF^low^–AXL^high^)^75^.

Based on these findings, we hypothesized that co-amplification of LZTR1 and CRKL might lead to increased rates of distant recurrence. Given that 22q11.21 amplification is a late-arising event in acral melanoma (Fig. 2c**, left**), we tested this hypothesis by examining acral melanoma patients diagnosed with stage II or III disease. Indeed, in patients for whom distant metastasis-free survival (DMFS) data were available, focal amplification of 22q11.21 was associated with earlier development of distant metastatic disease, with a median lead time of nearly 1 year (Fig. 6c).

Finally, we investigated whether overexpression of LZTR1 or CRKL release normal human melanocytes from their dependency on growth factors, a common phenotype of metastatic melanoma cells^76^. While normal human melanocytes retained their growth factor dependency (**Supplementary Fig. 8**), LZTR1, but not CRKL, enabled immortalized mouse melanocytes to form colonies and divide in the absence of their only required growth factor, TPA (tetradecanoyl phorbol acetate)^77^ (Fig. 6d,e). This phenotype is likely the consequence of MAPK activation, as seen by the presence of phosphorylated ERK (Fig. 6f).

Taken together, these results strongly implicate LZTR1 and CRKL in malignant transformation and the initiation of metastasis. While both genes showed similar phenotypes, the effects of overexpression were notably enhanced when LZTR1 and CRKL were co-expressed. However, only LZTR1 released immortalized mouse melanocytes from their dependency on growth factor, a characteristic shared by melanoma cells.

## DISCUSSION

Acral melanoma has high incidence among non-Caucasian populations, accounting for up to 86% of melanomas diagnosed in Asian patients as compared to ∼10% of Caucasians^29, 78–84^. Our work establishes common features of acral melanomas in cohorts from Asian and Caucasian populations. These include 1) the consistent association between specific focal amplifications and poor outcomes, and 2) the identification of *LZTR1* as a key gene within 22q11.21, the most prognostic recurrent alteration identified in both acral cohorts. Based on these findings, we performed a comprehensive analysis of LZTR1 signaling pathways and obtained functional evidence for LZTR1 as a tumor promoter.

LZTR1 is co-amplified with CRKL and downregulation of each gene inhibits melanoma cell proliferation, albeit to varying degrees. While CRKL has been linked to tumor growth as a candidate oncogene in several human malignancies, including lung adenocarcinoma^47–49^, LZTR1 is generally considered a tumor suppressor. Germline mutations in LZTR1 are involved in Noonan syndrome^53, 85^, schwannomatosis^46^ and glioblastoma^43, 86^. Moreover, somatic loss-of-function mutations in LZTR1 occur in 22% of glioblastomas. These mutations drive self-renewal and growth of glioma spheres^43^, consistent with a role in tumor suppression. However, despite these findings, LZTR1 is amplified in a subset of carcinomas (up to 8.3%), including bladder, uterine, and lung cancers^87, 88^. These data, coupled with our results, suggest that LZTR1 could have tumor-promoting capabilities in multiple human malignancies.

Unique aspects of our study include the broad range of tumor specimens analyzed and the utilization of cells harboring different oncogenes that modulate LZTR1 activity. For example, in *NRAS*-mutant melanoma, RAS mis-localized to the cytoplasm in response to shLZTR1 and caused MAPK inhibition. On the other hand, elimination of LZTR1 in BRAF-mutant cells increased BRAF levels, leading to ERK activation. In several cell lines, ERK activation induced growth arrest via MITF degradation, a process unique to melanocytes and the melanoma system^67, 68^.

Importantly, our study demonstrates that *LZTR1* and *CRKL* – two of the top three genes associated with ch22q11.2 amplification in acral melanoma – facilitate anchorage-independent growth in normal human melanocytes, likely by reducing E-cadherin, increasing N-cadherin, and activating integrin β1. The reciprocal expression of E-cadherin and N-cadherin in early melanoma progression has been known for about two decades^73, 89^, but to our knowledge, our findings link these events to genomic modification of two specific genes for the first time. In addition, we observed activation of MAPK and SRC kinases, the likely consequences of integrin signaling^90, 91^. The ability of LZTR1 to convert immortalized mouse melanocytes to a growth-factor independent mode of proliferation, a major characteristic of melanoma cells in culture, further underscores its tumorigenic potential. These results agree with our genomic observation that ch22q11.21 amplification is a late-arising event associated with regional and distant metastasis.

Separately, we identified a striking inverse relationship between immunomodulatory genes and 22q11.21 amplification. It is tempting to speculate that high levels of LZTR1 reduce the inflammatory response while protecting cells from stress-induced apoptosis, thereby facilitating metastasis. Conversely, patients with low levels of LZTR1 preferentially harbor a hot tumor microenvironment, which might provide benefit from immunotherapy. Future studies will be needed to explore these possibilities.

In summary, we demonstrate that late-arising focal amplifications of cytoband 22q11.21 are a leading determinant of shorter survival time in acral melanoma. Our genomic and functional experiments provide critical new insights into the pathogenesis of this disease and strongly implicate LZTR1 as a novel tumor promoter and promising therapeutic target.

## Acknowledgements

This work was supported by the Melanoma Research Alliance (R.H., Q.Y., J.C.), the Sokoloff-MRA Award (R.H.), the Yale SPORE in Skin Cancer (Bosenberg and Kluger), the Roslyn and Jeremy Meyer Award (R.H.), the National Cancer Institute (A.M.N., R00CA187192), the Stinehart-Reed foundation (A.M.N.), the Stanford Bio-X Interdisciplinary Initiatives Seed Grants Program (IIP) (A.M.N.), the Virginia and D.K. Ludwig Fund for Cancer Research (A.M.N.), the Natural Science Foundation of China Major Projects of International Cooperation and Exchanges grant 81620108024 (X.C.), the General Program grant 81874138 (M.Y.), a New Investigator Award provided by Rutgers Cancer Institute of New Jersey (State of NJ appropriation and National Institutes of Health grant P30CA072720, J.C.), and a Melanoma Research Foundation Career Development Award (J.C.). We wish to acknowledge the Yale Center for Genome Analysis (YCGA) for performing WES and RNA-seq, David Calderwood and Ben Turk for providing the short hairpin RNA lentiviral vectors, the West Campus Imaging Core for confocal microscopy and cell imaging, Junkun Liu for performing cell immunostaining, Robert Straub and Jenna Ollodart for technical assistance. We thank for Dr. Doug Brash for his insight regarding acral melanoma mutations and to Zoe Halaban for her critical questions and enthusiastic support during these studies. We dedicate this manuscript to the memory of our colleague Dr. Deepak Narayan, a surgeon-scientist who provided great insight into this work and who will be remembered for his legacy of unparalleled innovation in the face of complex problems.

## Author Contributions

A.M.N. and R.H. conceived the study, developed strategies for related experiments, and wrote the manuscript. X.C. initiated the collaborative studies, supervised sample collections from multi-clinical centers in China, exome and RNA-sequencing and some functional studies. F.F. co-wrote the paper and performed bioinformatics analyses with assistance from C.L., J.K., K.R., and L.P. R.H. performed key functional experiments with assistance from C.P., A.B., M.Y., M.Z., and J.N. A.B. performed experiments, tissue collection and processing (Yale cohort) and obtained clinical data. C.P. performed sample and clinical data collection and assisted with sequencing (CSU cohort). J.S. assisted with the collection of clinical data (CSU cohort). M.S. identified patients and collected clinical data (Yale cohort). S.A., D.N., J.C., K.O, S.Z., Y.W, G.H, M.W., and X.L contributed tumor specimens. J.C. and Q.Y. assisted with the conception of the study, performed experiments, and contributed to writing. All authors commented on the manuscript at all stages. A.M.N. and R.H. jointly supervised this work.

## Methods

### Human subjects

All clinical specimens in this study were collected with informed consent for research use and were approved by the Yale University and Central South University Institutional Review Boards in accordance with the Declaration of Helsinki. Melanoma tumor specimens were excised to alleviate tumor burden. CSU samples were collected from Xiangya Hospital, Hospital for Skin Diseases (Institute of Dermatology), Chinese Academy of Medical Sciences in Nanjin, Third Affiliated Hospital of Sun Yat-sen University, Hunan Provincial Tumor Hospital, Xiangya Hospital, Central South University, First Affiliated Hospital of Harbin Medical University and Wuhan Union Hospital.

### Nucleic acids extraction

Melanomas were sequenced from snap-frozen tumors (Yale and CSU cohorts) or low passage cell cultures (<4) as previously described^2, 4^. DNA from melanoma cells and freshly frozen tumors was extracted with the DNeasy purification kit (Qiagen Inc., Valencia, CA). High melanin content was removed with *OneStep*™ PCR Inhibitor Removal Kit (Zymo Research Corporation, Irvine, CA). Direct-zol™ RNA MiniPrep w/ Zymo-Spin™ IIC Columns (Zymo cat # D4019) were used to extract RNA from tumors, and the RNeasy PowerLyzer Tissue & Cells Kit (Qiagen, CAT # 15055-50) was used to extract RNA from peripheral blood mononuclear cells (PBMCs) and melanoma cells.

### Whole exome sequencing

#### Sample preparation

The quality of genomic DNA was determined by estimating the A_260_/A_280_ and A_260_/A_230_ ratios by nanodrop, both of which required to be >1.8, and by electrophoresis in 1% agarose gel in which high quality DNA migrates as a single high molecular weight band. One µg of genomic DNA was sheared to a mean fragment length of about 220 bp using focused acoustic energy (Covaris E220). The size distribution of the fragmented sample was determined by using the Caliper LabChip GX system. The fragmented DNA samples were transferred to a 96-well plate and library construction was completed using a liquid handling robot. Following fragmentation, we added T4 DNA polymerase and T4 polynucleotide kinase that blunt end and phosphorylate the fragments. The large Klenow fragment then adds a single adenine residue to the 3’ end of each fragment and custom adapters (IDT) are ligated using T4 DNA ligase. Magnetic AMPure XP beads (Beckman Coulter) were used to purify and size select the adapter-ligated DNA fragments. The adapter-ligated DNA fragments were then PCR amplified using custom-made primers (IDT). During PCR, a unique six base index was inserted at both ends of each DNA fragment. Sample concentration was determined by picogreen and the fragment length distribution using the Caliper LabChip GX system. Samples yielding at least 1 µg of amplified DNA were used for capture.

#### Targeted capture and sequencing

For the CSU cohort, capture was performed using the NimbleGenSeqCap Med Exome 44M kit, followed by 151 bp paired-end sequencing on the Illumina HiSeq X 10 platform. TrimGalore (version 0.3.7) and FastQC (v0.11.2) were used to remove adapters and low-quality sequences from the raw data. For the Yale cohort, equal amounts of each sample were pooled prior to capture. Example: for 16 samples per lane 62.5 ng of each genomic DNA library was pooled (1 µg total) and lyophilized with Cot-1 DNA and universal adapter blocking oligos (IDT). The dried sample was reconstituted according to the manufacturer’s protocol (IDT), heat-denatured, and mixed with biotinylated DNA probes produced by IDT (xGen Exome Panel). Hybridizations were performed at 65°C for 16 hours. Once the capture was complete, the samples were mixed with streptavidin-coated beads and washed with a series of stringent buffers to remove non-specifically bound DNA fragments. The captured fragments were PCR amplified and purified with AMPure XP beads. Samples were quantified by qRT-PCR using a commercially available kit (KAPA Biosystems) and insert size distribution determined with the LabChip GX. Samples with a yield of ≥0.5 ng/µl were used for sequencing. Sample concentrations were normalized to 2 nM and loaded onto Illumina NovaSeq 6000 flow cells at a concentration that yields at least 600Gbp of passing filter data per lane. Samples were sequenced using 101 bp paired-end sequencing reads according to Illumina protocols.

### Bulk and single-cell RNA sequencing

#### Bulk RNA-seq

For the CSU cohort, total RNA was depleted of rRNA using the Ribo-Zero rRNA removal kit, namely,1 μg of total RNA was used as input for rRNA removal. Sequencing libraries were generated using the TruSeq RNA sample prep kit (Illumina). The libraries were sequenced as 151 bp paired-end reads using an Illumina HiSeq X Ten platform. For the Yale cohort, rRNA was depleted starting from 25-1000ng of total RNA using the Kapa RNA HyperPrep Kit with RiboErase (KR1351). Indexed libraries that met appropriate cut-offs for both quantity and quality were quantified by qRT-PCR using a commercially available kit (KAPA Biosystems) and insert size distribution was determined with the LabChip GX or Agilent Bioanalyzer. Samples with a yield of ≥0.5 ng/ul were used for sequencing. Samples were run on a combination of Illumina HiSeq 2500, HiSeq 4000, and NovaSeq instruments, and multiplexed using unique dual barcode indexes (to avoid sample contamination or barcode hopping).

#### scRNA-seq

To obtain a single-cell transcriptional portrait of a chr22q11.21-amplified tumor, we analyzed a primary acral melanoma specimen (YUJASMIN, Yale cohort) with 6 focal copies of 22q11.21, as determined by WES. The 10x Chromium 5’ expression profiling platform with V1 chemistry was applied to a cryopreserved tumor cell suspension from YUJASMIN sorted for viable singlets to target 10,000 cells. Cells were sorted in the following ratios prior to library preparation: 50% CD3^+^CD45^+^ T cells: 25% CD3^−^CD45^+^ non-T immune cells: 25% CD45^−^ stromal/cancer cells. Cell viability was assessed by the LIVE/DEAD™ Fixable Red Dead Cell Stain Kit (catalog #L34971, Thermo Fisher). The following antibodies were used: Alexa Fluor® 488 anti-human CD45 antibody (clone H130, catalog #304019, BioLegend); APC anti-human CD3 antibody (clone HIT3a, catalog #300319, BioLegend). The 10x library was sequenced on an Illumina HiSeq 2500 instrument.

### Tumor genotyping from whole exome sequencing data

Sequencing reads from exome-captured samples were analyzed with a combination of germline and somatic variant calling, permitting the identification of somatic variants, loss-of-heterozygosity (LOH) regions and copy-number variation (CNV) regions.

BAM files of aligned reads were created for each sample by aligning the sequencing reads to the GRCh37 human reference with decoy sequences (the “hs37d5” reference) using BWA MEM^95^, marking duplicates using Picard MarkDuplicates (http://broadinstitute.github.io/picard), and then performing indel realignment and base quality score recalibration using GATK v3.2^96^. Then, variants were called using the tumor/normal bam files in three ways: 1) a joint variant call using GATK HaplotypeCaller, GenotypeGVCFs and hard filtering following GATK 3.2 best practices; 2) somatic SNP variant calls using MuTect with options “max_alt_alleles_in_normal_count=6”, “max_alt_allele_in_normal_fraction=0.1” and “max_alt_alleles_in_normal_qscore_sum=200”; 3) somatic indel variant calls using Indelocator with options “minCoverage=6”, “minNormalCoverage=4” and “minFraction=0.2”. The output from the three variant callers were merged using in-house scripts into a single VCF file, containing the union of GATK variants and MuTect/Indelocator somatic variants, marking variants called as somatic by MuTect or Indelocator as “somatic”.

Those variants were annotated using Annovar^97^ and VEP^98^, and then the somatic variants were filtered using the following criteria: 1) tumor alt depth ≥ 4, 2) normal read depth ≥ 4, 3) normal alt depth ≤ 1 or normal alt frequency less than 1/5 tumor alt frequency, 4) the maximum population frequency of the variant from ExAC^99^, NHLBI, 1000 Genomes, or Yale Exome database must be less than 2% for a cancer-related gene (any gene in the Oncomine or Foundation Medicine gene panels or COSMIC CG Census gene list) or 1% for any other gene. Also, only protein changing variants with a VEP impact of MEDIUM or HIGH, or variants within 15 bases of a protein coding splice site were reported in the final output.

Loss-of-heterozygosity (LOH) regions were identified using the joint variant calls generated from GATK^100^. For each variant that was called heterozygous in the normal and had a depth ≥ 20 in the normal, the allele frequency of the tumor and normal were subtracted (“abs (tumorAF – normalAF)”). Then the R loess and predict functions were used to smooth the allele frequency differences, and then any region with fitted values above 0.1 were identified as LOH. *Tumor Purity:* The tumor purity was estimated from variant allele frequency differences between tumor and normal, where tumor-specific allele differences (most commonly, homozygous alleles of the tumor in loss-of-heterozygosity (LOH) regions) shift the overall tumor sample’s allele frequencies proportional to the fraction of tumor cells in the sample. For each variant in the GATK joint calling, where the normal sample was called heterozygotic and had depth ≥ 20, the *allele frequency difference* (AFD) was calculated as “abs (tumorAF – normalAF)”), subtracting tumor from normal allele frequency. Those differences across the genome were smoothed using the R loess and predict (“predict(loess(AFD, span=0.28))”), and regions >= 0.1 were considered as allelic deviations from the normal. The mean of each region was calculated, and the maximum mean was doubled to provide the tumor purity estimate (as, for example in an LOH region, the deviation from normal heterozygotic calls is one-half the tumor fraction, plus or minus one of the two tumor alleles).

Copy-number variant (CNV) regions were identified by first calculating the mean read depth for each RefGene coding exon, for the tumor and normal samples. Normalized tumor/normal read depth ratios were computed for each exon (normalized by the mean read depth of the tumor and normal across the exome), and then, using a partitioning of the genome into 20kb bins, a mean ratio for each 20kb region of the genome, which contains an exon, was computed. Those ratios were converted to log-ratios, then de-noised and segmented by circular binary segmentation (CBS) using the DNAcopy library from R (http://bioconductor.org/packages/release/bioc/html/DNAcopy.html.) Regions with a value deviating from 0.0 are identified as CNV’s. For each CNV, the log-2 ratio is converted back to a simple ratio and the copy number is calculated as “2 + round((ratio – 1.0) / step))”, where “step” is one of three values based on the estimated tumor purity, specifically 0.5, 0.4 or 0.32 for samples with purity ≥ 80%, purity between 40% and 80% or purity <= 40%, respectively.

Focal amplifications and deletions were identified with GISTIC2.0 (version 2.0.23, release date 27 Mar 2017)^33^ using the CBS segmentation files described above and the hg19 reference genome (GRCh37). No marker input file was provided. Parameters were specified according to the authors’ recommended run profile: amplification and deletion thresholds were set to 0.1, the q value threshold was set to 0.1 with a confidence level of 0.95, and log_2_ ratios were capped at 1.5. Gene-level GISTIC analysis and broad analysis were also applied, with a focal length cutoff of 0.7. Wide peaks identified with a *Q* value less than 0.1 in each melanoma subtype were aggregated into a master list, and genes within each wide peak were used to construct a copy number matrix. Of note, if two or more peaks were identified within the same cytoband, we appended a suffix to the cytoband name to denote the melanoma subtype in which the cytoband was identified (AC, acral; SE, sun-exposed). If more than one peak was identified within the same cytoband for a given melanoma subtype, the subtype acronym was followed by a numerical identifier (1, 2, etc.). For cases in which one peak completely encompassed another one and where both peaks had the same orientation (i.e., amplified or deleted), the shorter one was eliminated.

For comparative genomics and survival analyses, we constructed a matrix containing the mean copy number per wide peak (rows) for each melanoma tumor sample (columns). The mean copy number per wide peak was calculated as the average gene-level copy number (estimated as described above) per wide peak. We subtracted 2 from all gene-level values so that copy number-neutral regions are equal to 0.

MAF files for each sample’s somatic variants were generated by vcf2maf (version 1.6.14), using Variant Effect Predictor release 91 and filtering by ExAC version 0.3. The acral variants, and separately the sun-exposed variants, were combined into a single MAF file and given to MutSigCV (version 1.41)^101^, which was run using default parameters and the exome coverage, covariate and dictionary files included in the MutSigCV package.

### Visualization of somatic alterations across patients

Recurrently mutated genes and significant focal CNVs were visualized using the *Oncoprint* function in *ComplexHeatmap*^102^. The default bar plot (top) was replaced with a bar plot showing the number of nonsynonymous SNVs and indels per patient. For CNV regions, amplifications and deletions were calculated by averaging the GISTIC-generated gene-level copy numbers for all genes within each wide peak. Peaks with an average copy number above 3 (one additional copy) were deemed amplified, whereas peaks with an average copy number below 1.4 were considered deleted.

### Bulk RNA-seq analysis

Raw RNA-seq reads were aligned with Salmon^103^ (version 0.99) to the GENCODE v.25^104^ reference transcript assembly. Subsequently, the tximport^105^ was used to generate an expression matrix normalized to transcripts per million (TPM). Protein-coding genes were determined using Ensembl release 92 human annotation^106^ (GRCh38.p12, Apr 2018), extracted by biomaRt^107^ (version 2.40.5) and non-protein-coding genes were omitted. Expression values were renormalized to TPM after this step. For batch normalization, we applied *ComBat* from *sva*^108^ using a parametric adjustment for sequencing center and year of sequencing. Following batch correction, negative values were replaced with zero and the expression matrix was log_2_-transformed after adding a pseudo-count of 1.

To delineate pathways associated with focal amplification of 22q11.21 (Fig. 2d), genes differentially expressed between 22q11.21 amplified and non-amplified acral melanomas were identified by constructing a linear model (*lm* function in R) to predict amplification status as a function of 1) gene expression (log_2_ adjusted) and 2) tumor stage (at the time of resection). The *t* value corresponding to the expression vector of each gene was used to rank-order the transcriptome. Pre-ranked gene set enrichment analysis (GSEA)^109^ was subsequently applied to the ranked-ordered transcriptome in order to assess HALLMARK pathways in MSigDB (version 7.2)^110^.

Related to Fig. 2e, we curated a list of immunomodulatory genes, including immune checkpoint molecules, and analyzed their expression in both acral cohorts using hierarchical clustering applied with Pearson correlation and Ward D2.

### Gene fusion detection from RNA-seq data

To identify fusion genes, we aligned the RNA-seq reads for each sample to the GRCh38 human reference genome using HISAT2^111^. Candidate fusion transcripts in the sequencing reads were identified with STAR-Fusion, employing the STAR aligner and FusionInspector annotator to identify the position of the chimeric RNA. For ease of manual review, the fusions were sub-grouped, with each fusion placed into the first group that either gene matched: 1) mitochondrial genes, 2) immunoglobulin genes, 3) protocadherin genes, 4) commonly expressed fusions using GTEx expression data, 5) fusions of neighboring/local_rearrangement genes, 6) non-annotated genes, and 7) all others i.e., the rare, non-local fusions of annotated protein coding genes.

### Survival analysis

Cox proportional hazards regression was applied to estimate overall survival associations. Cases with an initial diagnosis preceding the sequenced tumor by more than 5 years were excluded from analysis (*n* = 5). To estimate stage-adjusted associations with overall survival, stage was added as a covariate. Kaplan-Meier plots for comparison of survival curves were generated either by the *survminer* ^112^ package in R (version 0.4.5) or by Graphpad Prism (version 8).

To determine survival associations of focal CNVs identified by GISTIC (Fig. 2a, **Supplementary Fig. 3a**), we applied Cox regression separately to each region within each acral melanoma cohort using the copy number matrix described above (see *Copy number analysis*). In all cases, we dichotomized each CNV by analyzing amplified (>0) versus non-amplified (≤0) and deleted (<0) versus non-deleted (≥0). Survival z-scores were combined across cohorts using Stouffer’s method^113^, yielding an unweighted meta-z-score for each gene.

To analyze SNV- and indel-related survival associations in acral melanomas, we examined genes harboring one or more nonsynonymous mutations with at least 5% recurrence frequency in either acral melanoma cohort. These data were used to create a binary matrix in which recurrently mutated genes were rows and patients were columns (1, at least one recurrent SNV or indel; 0, otherwise). Survival z-scores were combined across cohorts as indicated above. The following four genes were insufficiently recurrent in at least one cohort to run Cox regression: *CGREF1*, *NF1*, *TP53*, and *ARID2*. To calculate survival associations for these genes, we randomly up-sampled patients from the Yale cohort in order to match the size of the CSU cohort. We then generated a cross-cohort survival *Z*-score for each gene.

To relate *LZTR1* expression to overall survival, we dichotomized patients in each cohort by determining an expression threshold that discriminates 22q11.21-amplified from non-amplified patients at a defined specificity. This was done to link the threshold for dichotomization with 22q11.21 amplification without being confounded by the upper range of *LZTR1* expression in non-amplified tumors. We used a specificity cutpoint of 95% for acral melanomas profiled in this study and sun-exposed melanomas profiled by TCGA. A specificity cutpoint of 90% was used for sun-exposed melanomas profiled in this work owing to a lack of evaluable samples in the ‘high’ group (n=1) at a specificity cutpoint of 95%. Notably, *LZTR1* expression was also significantly associated with adverse outcomes when assessed as a continuous variable (i.e., without dichotomization) in sun-exposed melanomas profiled by TCGA (*Z* = 2.86, *P* = 0.004). Skin cutaneous melanoma (SKCM) expression, copy number, and survival data from TCGA were downloaded from cBioPortal^87^.

### Single-cell RNA sequencing analysis

Single-cell RNA-seq reads were mapped to the GRCh38 human reference assembly and barcode-deduplicated using Cell Ranger (version 3.0.2). In total, 7,551 cells were sequenced, yielding a median of 945 genes per cell. The expression matrix generated by Cell Ranger was converted to counts per million (CPM) and was log_2_ transformed after addition of a pseudo-count to every value. Cells with less than 500 expressed genes were removed before importing the data into *Seurat* ^114^ (version 3.0.2). Seurat was applied for pre-processing (default settings), data normalization (default settings), identifying the most variable features, dimension reduction (PCA and UMAP), finding marker genes, and clustering the single-cell expression data. In finding variable features, the low and high mean cutoffs were set to 0.0125 and 3, respectively, and the dispersion cutoffs were respectively set to 0.5 and infinity, with 20 bins. PCA was generated on the most variable genes with the first 15 principal components. Neighbors were calculated from the first 13 PCA components (as determined by the JackStraw function), followed by cluster analysis (resolution = 0.5).

#### Single-cell copy number analysis

To estimate large-scale copy number alterations from scRNA-seq data, we used CONICSmat (Copy-Number analysis In single-Cell RNA-Sequencing from an expression matrix)^36^ as implemented in the R package, *CONICSmat*. Per the authors’ recommendations, raw read counts were divided by 10 and were log_2_-adjusted after adding one to each count value. Chromosome arm positions from GRCh38 were used to define arm coordinates. Briefly, genes expressed in 5 cells or less were filtered, a normalization factor for each cell was calculated, and a Gaussian mixture model was calculated based on the z-score of the average centered gene expression for each region across all cells. Melanoma cells (*n* = 321) were split into two groups based on the estimated relative copy number in the 22q arm. Cells with a normalized copy number between 0 and 0.10 were used for normalization of the expression data. Histograms were generated using *plotHistogram* and by setting the z-score threshold to 4. Chromosomal alterations in each single cell were visualized by *plotHistogramHeatmap* with the authors’ recommended parameters (window size = 120, expression threshold = 0.2, visualization threshold = 1). Of 12 genes identified by GISTIC within the chr22q11.21 wide peak, relative copies of 22q11.21 for each cell were calculated by averaging the estimated copy number for all 7 genes with detectable expression (CPM>0).

To characterize the RNA expression profile associated with chr22q11.21 focal amplification, the median relative copy number (0.45) inferred by CONICSmat was used to split the cells into chr22q11.21 high and low groups. The log_2_-adjusted scRNA-seq dataset normalized by Census ^115^ was compared between these two groups to identify differentially expressed genes using a two sided *t*-test with unequal variance, and the resulting *t*-statistics were used for ranking the gene list. This gene list was submitted to pre-ranked GSEA^109^ to interrogate 50 HALLMARK pathways (1,000 permutations, weighted enrichment statistics, MSigDB version 7.2^110^).

#### Single-cell differentiation status

To predict the relative differentiation status of each melanoma cell profiled by scRNA-seq, we used CytoTRACE^42^, a computational framework for inferring developmental potential on the basis of transcriptional diversity. Each of the 321 acral melanoma cells from chr22q11.21 amplified tissue received a CytoTRACE score between 0 (more differentiated) and 1 (less differentiated). CytoTRACE scores were visualized by Uniform Manifold Approximation and Projection (UMAP). To determine whether CytoTRACE is influenced by genes located on 22q, we reran CytoTRACE after excluding all genes on the 22q arm (**Supplementary Fig. 4e**).

### Short hairpin RNA (shRNA), CRISPR-Cas9 sgRNA, and cell viability tests

We used puromycin-bearing MISSION lentiviral vectors pLKO.1 shRNA to test the effect of downregulation of target proteins on cell proliferation and signal transduction, employing scramble vector SH002 as a negative control (MISSION, Sigma-Aldrich), or scrambled RNA. LentiCRISPRv2 plasmid was obtained from Addgene (addgene.org). Guide sequences targeting LZTR1 were designed using CHOPCHOP (https://chopchop.cbu.uib.no/) and cloned into LentiCRISPRv2 to generate single sgRNA carrying plasmids following a standard method^116^. A non-target sequence was included as the control.

The plasmids were packaged in lentiviral vectors with ViraPower™ Lentiviral Packaging Mix kit (Thermo Fisher, cat # K497500), and transfected into 293T cells. The medium was collected and filtered with Millex-GV 33 mm PVDF filter (Millipore SLGV033RS) and then concentrated with Amicon Ultra-15 centrifugal filters (Millipore UFC910024). Melanoma cells and normal human melanocytes were infected with the lentiviruses, medium was changed the following day, and the cells were then incubated with puromycin (2.5 µg/ml) for five days. Cells were collected and processed for western blotting with antibodies to target proteins. In addition, two days after infection the shRNA treated cells were seeded in 96-well plates in triplicate or quadruplet wells and tested for cell viability in the absence and presence of puromycin for 72 hrs with the CellTiter-Glo® Luminescent Cell Viability Assay, for apoptosis or RAS activity GTPase assay.

Alternatively, GV298-U6-MCS-Ubiquitin-Cherry-IRES-puromycin lentiviral plasmids were purchased from GeneChem, China. The plasmids were co-transfected with packaging plasmids (pspAX2 and pMD2G) into 293T cells using Turbofect (Thermo Scientific) according to the manufacturer’s instructions. Lentiviruses were collected after 48 and 72 hours and used to infect into acral melanoma cells. Infected cells were incubated in medium supplemented with puromycin (1 μg/ml), for two or three days, seeded in 96-well plates (2×10^3^/well, five replicates) and cell viability was measured with Cell Counting Kit-8 (CCK-8) (Bimake.com, China). The CCK-8 test was repeated every 24 hrs for three days.

### CRKL and LZTR1 lentivirus vectors

pDONR223-CRKL and pDONR223-*LZTR1* were purchased from Addgene and DNASU, respectively. *LZTR*1 was transferred into pInducer20 vector^117^ (a gift from Dr. Thomas F. Westbrook, Baylor College of Medicine), and CRKL to PLX304 with the Gateway LR Clonase II Enzyme mix (Thermo Fisher 11791020). The identity of each vector was validated by targeted sequencing. Lentiviruses were produced as described above and the pInducer20 LZTR1 infected melanocytes were selected with 250 mg/ml Geneticin (G418) from American Bio (Canton, MA), CAT # AB05057-05000), and PLX304CRKL with 2.5 µg/ml Blasticidin S HCl from Thermo Fisher Scientific (Waltham, MA), CAT # A1113903.

### Cell proliferation and apoptosis

The melanoma cells were grown in OptiMEM (Invitrogen, Carlsbad, CA) supplemented with 5% fetal calf serum and antibiotics. Normal human melanocytes (NBMEL) were grown from newborn foreskins in medium supplemented with bFGF, heparin, IBMX and dbcAMP^76^. Mouse melanocytes were grown from one-day old newborn pups in the presence of horse serum, TPA, melanotropin, isobutyl methyl xanthine, and placental extract. They became immortalized and were shifted to medium containing only TPA after ∼20 passages in cultures^118^. Some of the Yale melanoma cell lines were characterized by next-generation sequencing before^2, 4^.

Cell proliferation was measured with the CellTiter-Glo® Luminescent Cell Viability Assay (Promega Corporation, Madison, WI). Melanoma cells were seeded in 96-well plates in triplicate or quadruplet wells after knockdown with hairpin lentivirus shRNA as indicated. Standard Error (SE) was calculated employing GraphPad Prism 7 software^119^. In addition, we seeded cells in 12-well plates (10-15,000/well) and measured proliferation by counting the number of cells from triplicate wells over a period up to 7-9 days with Beckman Cell Counter. For cell count by crystal violet, we seeded cells (3×10^3^/well) in 6-well plates and then incubated for 10 days. Following incubation, cells were immobilized with 4% paraformaldehyde in PBS for 15 min, stained with crystal violet for 10 min and then washed with PBS. A minimum of three random fields at 40X magnification were counted to determine cell numbers. Each sample had three replicates.

The rate of apoptosis was measured using the Dead Cell Apoptosis Kit with Alexa Fluor® 488 annexin V and propidium iodide (Invitrogen, V13241) following manufacturer instructions.

PLX4032 (500 nM, Plexxikon)^119^ or LY3009120 (100 nM, Selleck, Pittsburgh, PA, Catalog No.S7842) were added to the growth medium four hours before harvesting the cells for Western blotting.

For 3D cultures, melanocytes were suspended in 1 ml medium and seeded on 0.5% collagen (Cultrex, R&D Systems, Minneapolis, MN, Cat # 3442-050-01), in 24 well plates for three days.

### Microscopy

Images were acquired using an inverted Nikon Eclipse Ti fluorescence microscope with a Plan Apochromat lambda 60X /1.40 Oil objective or a Plan Fluor 4X/0.13 objective for fluorescent images or DIC images, respectively, a CSU-W1 confocal spinning disk unit, an iXon Ultra 888 camera (Andor Technology), MLC 400B laser unit (Agilent Technologies) and NIS Elements software (Nikon).

### Western blotting and antibodies

We used western blots to identify the levels of proteins as previously described ^69^. Cell extracts (20 µg/lane) were fractionated in 3%-8% or 4-12 % tris-acetate gel (NP0006, NuPAGE Life Technologies). All antibodies were used at the concentrations recommended by the manufacturers.

### RAS activity assay

The amount of GTP-bound RAS was determined using the Ras GTPase Chemi ELISA Kit (Active Motif North America, 1914 Palomar Oaks Way, Carlsbad, C 92008) following the manufacturer’s protocol. Melanoma cells treated with control shRNA or shLZTR1 were collected five days after infection by scraping on ice, washed with cold PBS, lysed, centrifuged and 50 µg protein/assay, in triplicates, were used following the manufacturer instructions.

### Immunostaining

Cells were grown on the surface of 4-well slides, washed 2-3 times with PBS, fixed with 4% paraformaldehyde for 15 min at room temperature, washed three times with PBS, permeabilize with 0.2% NP40 in PBS for 5 minutes, washed with PBS and incubate in PBS containing 1% BSA or (blocking buffer) for one hour. The cells were incubated with anti-GM130 antibody (clone 4A3 Millipore, Mouse), or calnexin (mouse mAb) for 1 hr at room temperature, and stained with secondary Alexa Fluor (Cy2) diluted in blocking buffer 1:1000 for 1 hr. They were washed 3X with PBS, incubated with rhodamine-phalloidin to stain actin and DAPI to stain the nucleus.

### Statistical analysis

Linear relationships were modeled by linear regression (*R*^2^), and a *t* test was used to assess whether the result was significantly nonzero. When data were normally distributed, group comparisons were determined using a *t* test with unequal variance or a paired *t* test, as appropriate; otherwise, a Wilcoxon test was applied. Results with *P* < 0.05 were considered significant. Data analyses were performed with R and Prism v7 (GraphPad Software, Inc.). The investigators were not blinded to allocation during experiments and outcome assessment. No sample-size estimates were performed to ensure adequate power to detect a pre-specified effect size.

## Data availability

Raw sequencing data will be deposited in the Sequence Read Archive (SRA) and https://www.biosino.org/node/ and expression data will be deposited in the Gene Expression Omnibus (GEO). Accession numbers are pending.

