## Supplementary Figures for "Integrative molecular and clinical profiling of acral melanoma identifies LZTR1 as a key tumor promoter and therapeutic target"

**Table of Contents:**

**Supplementary Figures**


| | |
|---|---|
| Supplementary Figure 1 | Characteristics and overall survival patterns of patients analyzed in this study |
| Supplementary Figure 2 | Landscape of focal amplifications and deletions in acral melanoma |
| Supplementary Figure 3 | Associations of 22q11.21 amplification status with survival, stage, and lymph node status in acral melanoma |
| Supplementary Figure 4 | Analysis of a 22q11.21-amplified acral melanoma tumor by scRNA-seq |
| Supplementary Figure 5 | Prognostic associations of *LZTR1* expression in melanoma |
| Supplementary Figure 6 | Impact of downregulating chr22q11.21 genes on cell proliferation |
| Supplementary Figure 7 | Analysis of cell cycle proteins in response to shLZTR1 |
| Supplementary Figure 8 | LZTR1 and CRKL, alone and in combination, failed to release human melanocytes from their dependency on growth factors |

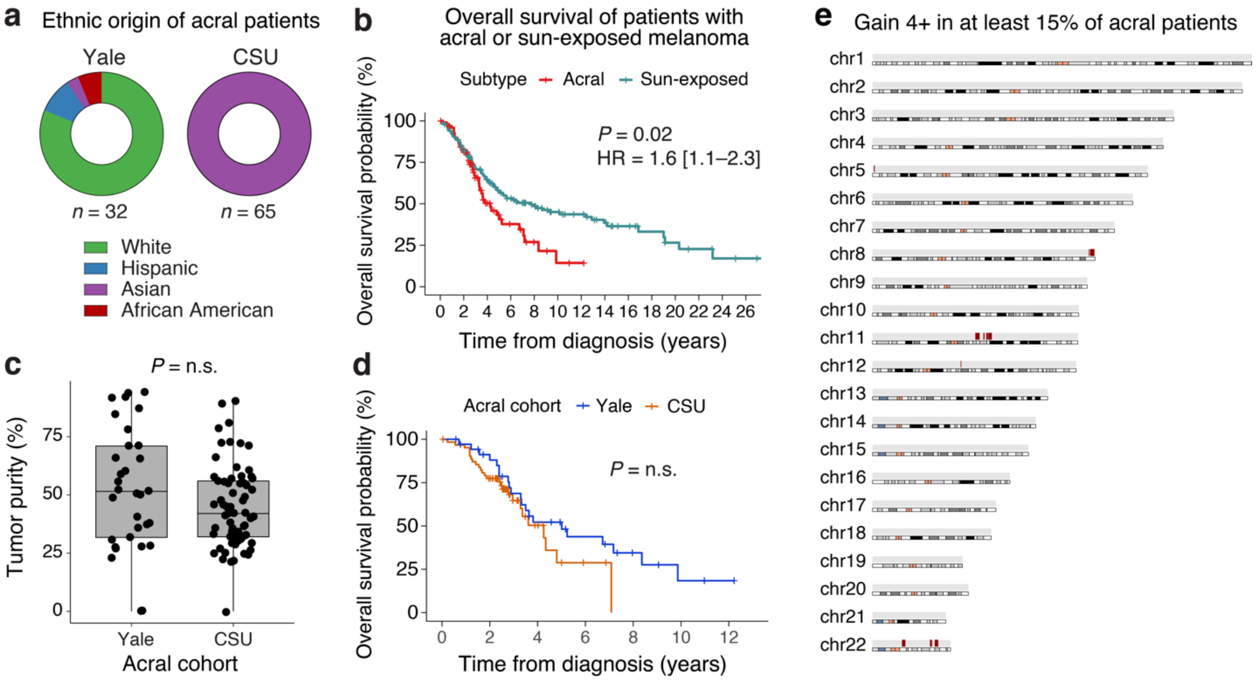


**Supplementary Figure 1: Characteristics and overall survival patterns of patients analyzed in this study. a**, Pie charts depicting ethnic origin of acral melanoma patients in each cohort (Table 1). **b**, Kaplan-Meier plot showing overall survival in acral versus sun-exposed melanoma subtypes analyzed in this work. Statistical significance was calculated with a log-rank test. HR, hazard ratio; HR 95% confidence interval is shown in brackets. **c**, Tumor purity of acral melanoma samples profiled by whole exome sequencing in Yale (*n* = 32) and CSU (*n* = 65) cohorts (**Methods**). Group comparison was performed by a two-sided Wilcoxon rank sum test. **d**, Kaplan-Meier plot showing overall survival of acral melanoma patients in Yale versus CSU cohorts. Statistical significance was calculated with a logrank test. n.s., not significant. **e**, Karyoplot of recurrent amplifications in acral melanoma. Chromosomal location of genes with a gain (red), defined as CNV ≥ 4 in ≥ 15% of acral melanomas analyzed in this work demonstrate specific patterning of concentrated amplification events.

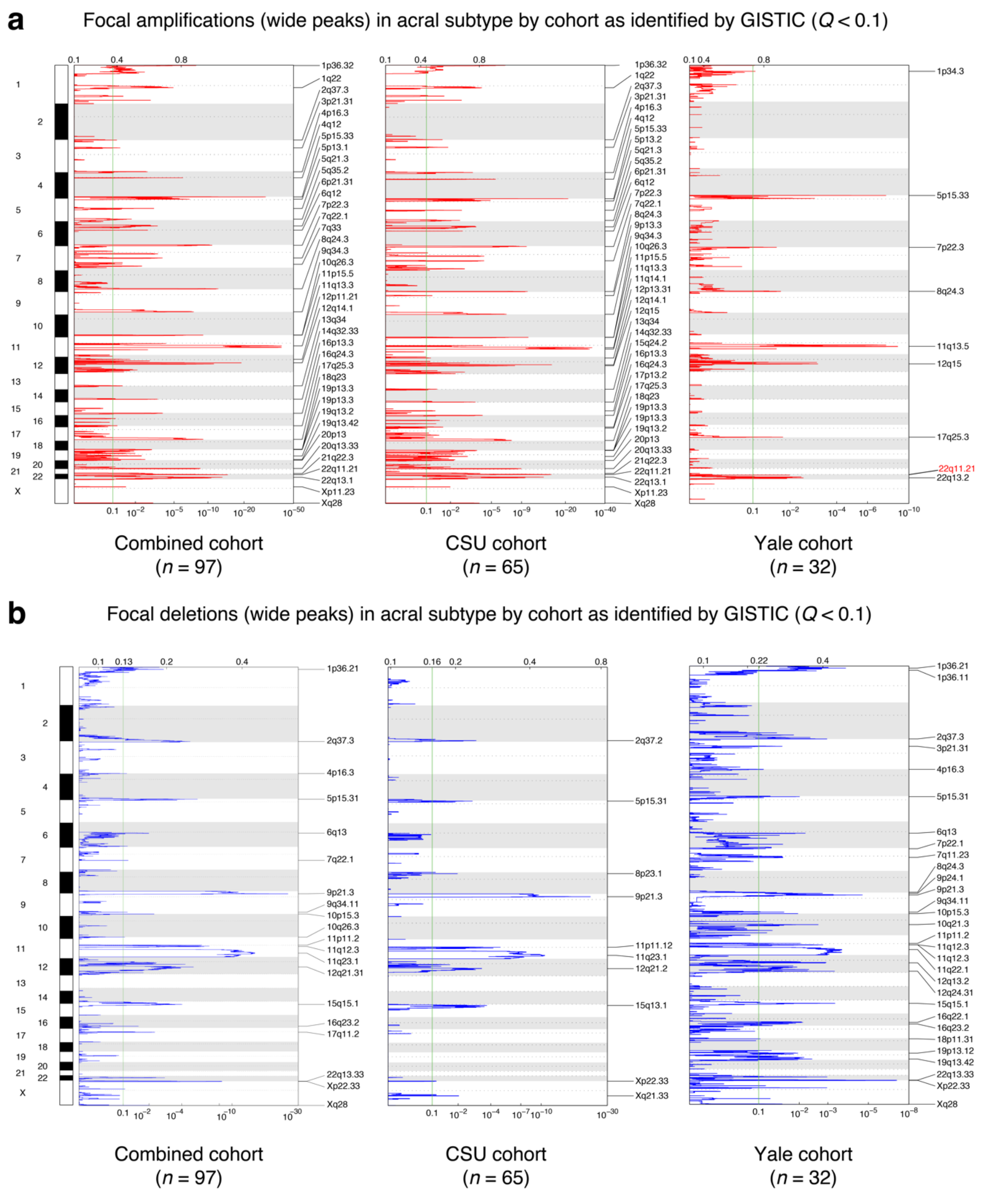


**Supplementary Figure 2: Landscape of focal amplifications and deletions in acral melanoma**. **a**, Focally-amplified wide peaks in acral melanoma tumors identified by Genomic Identification of Significant Targets in Cancer (GISTIC) (*Left*, all tumors; *Center*, CSU cohort; *Right*, Yale cohort). Labeled cytobands denote wide peaks with $Q < 0.1$. Although significant, 22q11.21 was not automatically labeled by GISTIC in the Yale cohort and is therefore indicated in red text. **b**, Same as a but showing focally-deleted wide peaks.

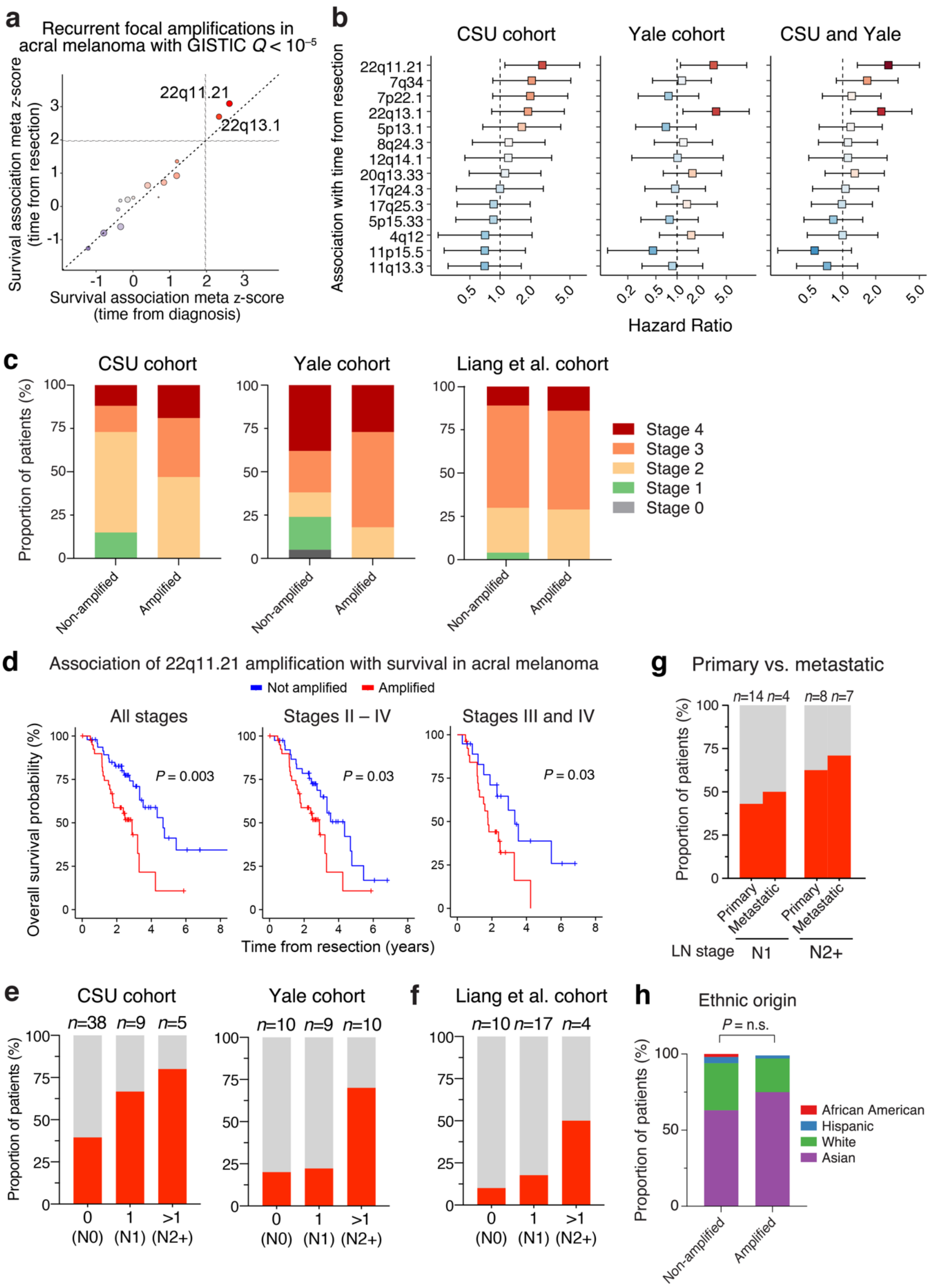


**Supplementary Figure 3: Associations of 22q11.21 amplification status with survival, stage, and lymph node status in acral melanoma. a**, Same as Fig. 2a, but showing only recurrent focal amplifications identified in acral melanomas (CSU and Yale combined) with GISTIC $Q < 10^{-5}$. **b**, Association between significant focal amplifications defined in the CSU cohort (GISTIC $Q < 10^{-5}$) and overall survival (from date of tumor resection), shown as hazard ratios (boxes) with 95% confidence intervals (error bars). Red indicates hazard ratio (HR) > 1 (worse prognosis) and blue indicates HR < 1 (better prognosis). Notably, while HRs are shown for CSU, Yale, and CSU + Yale (combined), focal amplifications in this analysis were defined from the CSU cohort alone. **c**, Stage distribution in patients with acral melanoma, shown as a function of 22q11.21 focal amplification status in three independent cohorts (CSU, Yale, Liang et al.). **d**, Association of chr22q11.21 amplification with overall survival in acral melanoma (CSU and Yale cohorts), shown for all stages, stage II to IV disease, and advanced disease (stage III and IV). Statistical significance was assessed by a log-rank test. **e**, Frequency of 22q11.21-amplified acral melanoma patients shown as a function of the number of positive lymph nodes. **f**, Same as **e** but for patients from Liang et al. **g**, Frequency of 22q11.21-amplilfied acral melanoma patients shown as a function of lymph node (LN) stage and stratified by tumor site of origin (primary or metastatic). **h**, Ethnic distribution of acral melanoma patients shown as a function of 22q11.21 focal amplification status. Statistical significance was assessed by a Chi-square test.

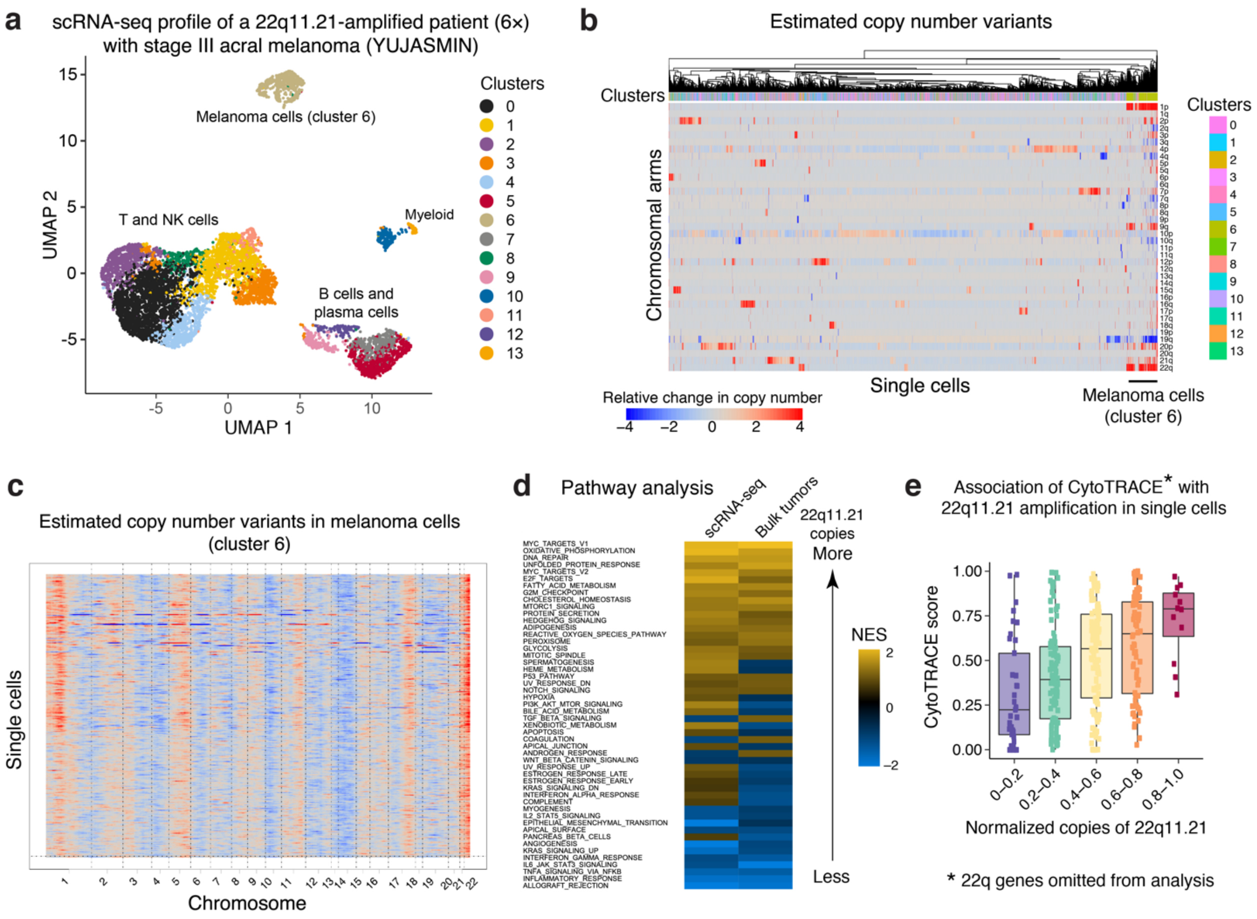


**Supplementary Figure 4: Analysis of a 22q11.21-amplified acral melanoma tumor by scRNA-seq. a**, UMAP projection of 7,166 single-cell transcriptomes from a 22q11.21-amplified patient (six copies) with stage III acral melanoma (YUJASMIN). Major cell lineages are annotated. Clusters were identified with Seurat (**Methods**). **b**, Heat map depicting arm-level copy number alterations estimated by CONICSmat in the cells from panel **a**, with columns ordered by hierarchical clustering. Each column represents a cell ($n$ = 7,166) and each row represents a unique chromosomal arm. Estimated amplifications of 22q and 1p are enriched in cluster 6 (melanoma cells, far right). **c**, Heat map portraying estimated copy number variants in melanoma cells ($n$ = 321; cluster 6) from panel a. Each row represents a cell. Columns (genes) are ordered by each gene's physical location in the genome. A potential subclone harboring amplifications in 1p and 22q is evident (top). Within this subclone, gene expression on the 22q arm is elevated, indicating single-cell variability in the copy number status of 22q. **d**, Pre-ranked gene set enrichment analysis (GSEA) showing concordance in hallmark pathways among melanoma single-cell transcriptomes from panel a (scRNA-seq) and 38 bulk acral melanoma tumors (related to Fig. 2d), in relation to high vs. low 22q11.21 copy number status (**Methods**). The former was stratified into high and low groups using the estimated relative copy number of genes within chr22q11.21 (**Methods**). The latter was calculated as in Fig. 2d. **e**, Same as Fig. 2g (bottom) but with genes on 22q omitted from the CytoTRACE analysis.

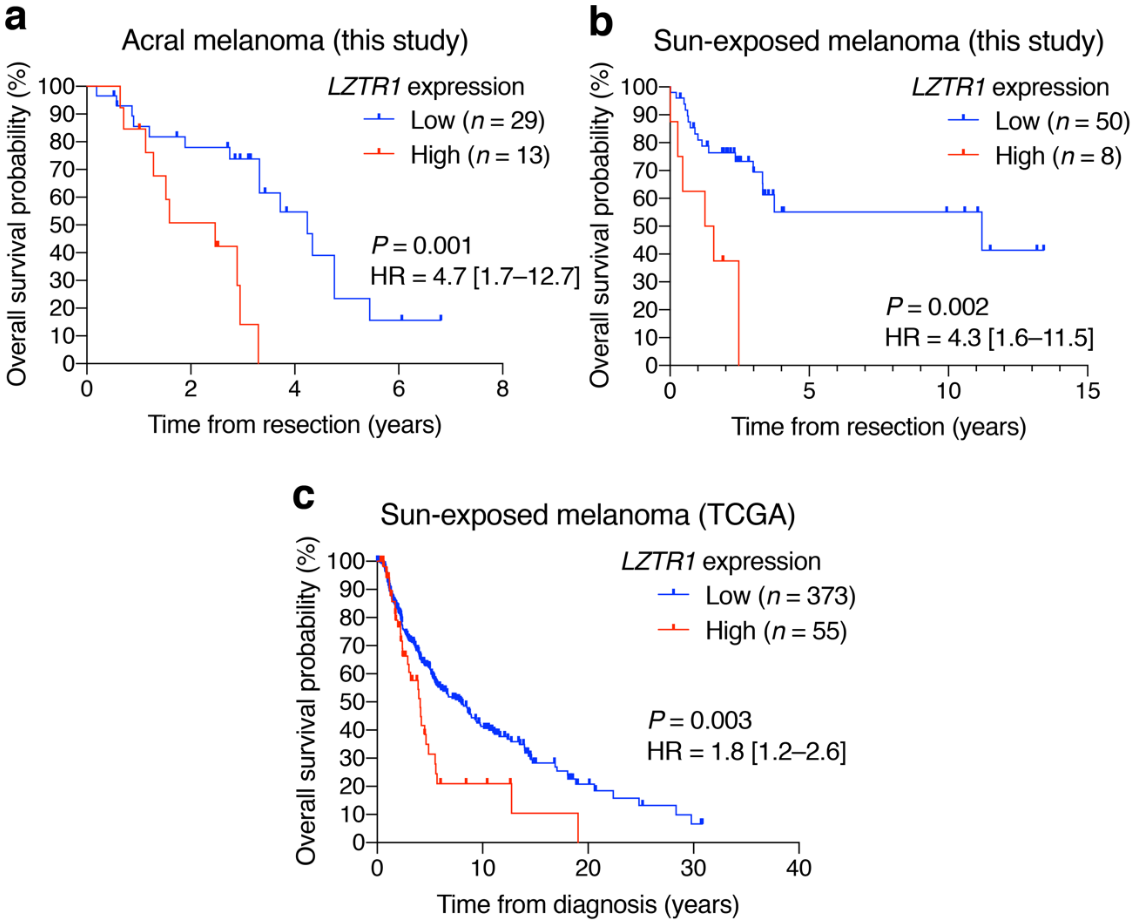


**Supplementary Figure 5: Prognostic associations of *LZTR1* expression in melanoma. a**–**c**, Kaplan-Meier plots showing differences in overall survival between LZTR1 high and low patients with acral and sun-exposed melanomas profiled in this work (panels a and b, respectively) and sun-exposed melanomas profiled by TCGA (panel c). LZTR1 was stratified into high and low groups on the basis of copy number status, as described in Methods. Statistical significance was determined by a log-rank test. HR, hazard ratio. 95% HR is indicated in brackets.

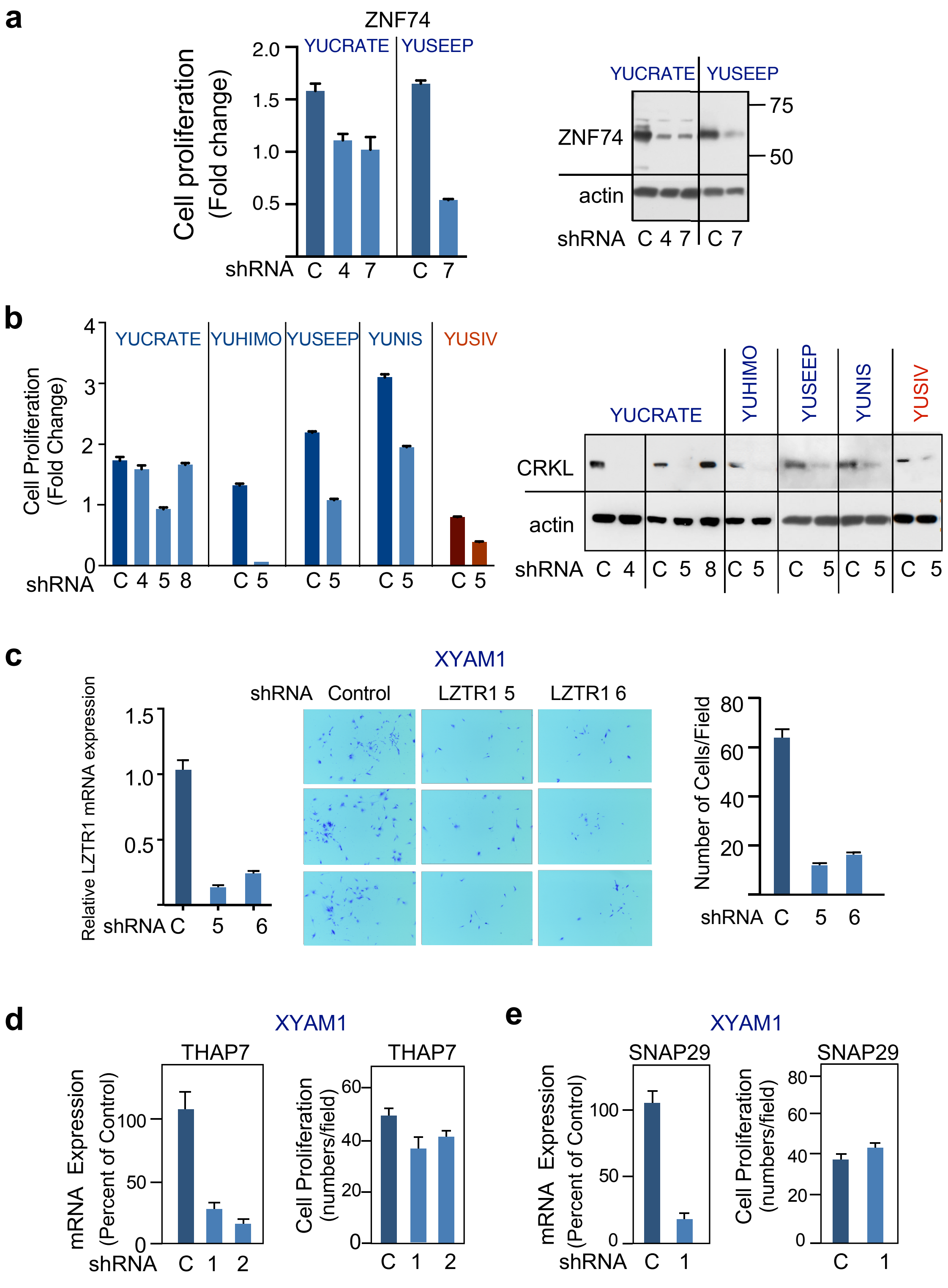


**Supplementary Figure 6: Impact of downregulating chr22q11.21 genes on cell proliferation**. **a**, Effect of ZNF74 shRNA on proliferation of primary acral melanoma cell lines (YUCRATE and YUSEEP) (left). Efficiency of ZNF74 knockdown (right). **b**, Effect of CRKL shRNAs on cell proliferation (left) and protein expression (right). Cell proliferation in a and b was assessed by a CellTiter-Glo® assay. Bar plots depict fold change between the 3rd and 6th day after infection with ZNF74 or CRKL shRNA (numbered), as compared to control (scrambled) shRNA (‘C’). Values are average of three or four readings ± SEM. **c**, Impact of LZTR1 knockdown on proliferation of XYAM1 acral melanoma cells. Shown are gene expression measured with RT-qPCR (left), representative images (middle), and quantification of clonogenic assays (right). XYAM1 acral melanoma cells were established from a patient treated in XiangYa Medical School, Changsha, China. Bar Plots on the left and right panels are means of triplicates ± SEM for LZTR1 shRNA 5 and shRNA 6, respectively. **d**, **e,** Knockdown of THAP7 (**d**) and SNAP29 (**e**) had little to no effect on proliferation of XYAM1 acral melanoma cells. Shown are gene expression measured with RT-qPCR (left) and quantification of clonogenic assays (right). Bar plots show means of triplicates ± SEM. Cell lines identifiers are indicated above all plots and colored according to their origin: acral melanoma (blue), sun-exposed melanoma (red).

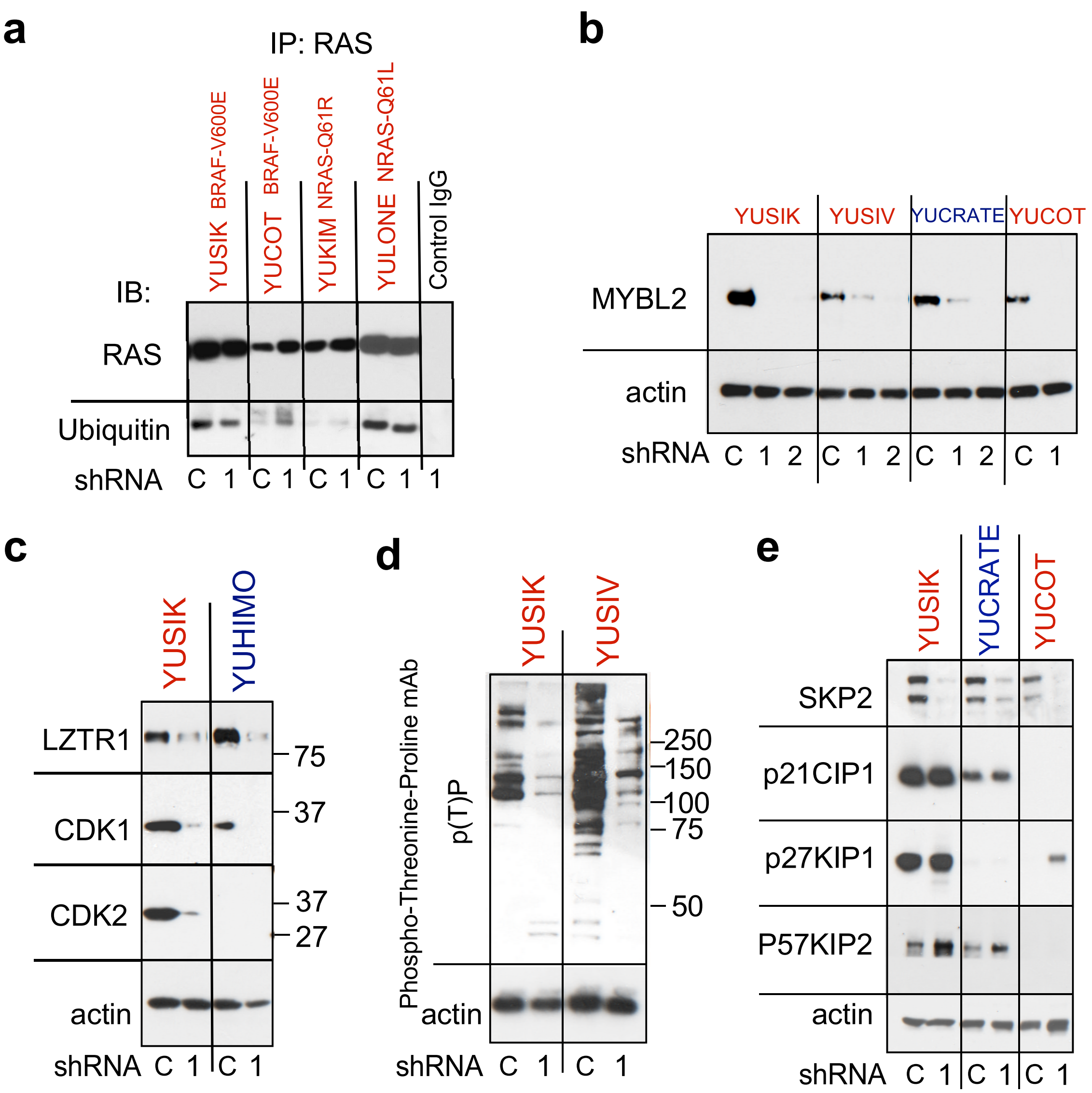


**Supplementary Figure 7: Analysis of cell cycle proteins in response to shLZTR1.** **a,** RAS ubiquitination is not altered in response to LZTR1 knockdown. Four independent cell lines were treated with shRNA scramble control ('C') or shLZTR1 ('1') for 6 days and cell extracts were used to immunoprecipitate RAS (IP) with anti-RAS (mouse monoclonal antibody) and immunoblot with anti-RAS or anti-ubiquitin, rabbit polyclonal antibodies (IB). Immunoprecipitation with IgG using cell extract from YUSIK was used as a control. **b**, **c**, Downregulation of cell cycle proteins predicted from GSEA analysis (**Fig. 5a**). Western blots showing downregulation of MYBL2, CDK1 and CDK2 in response to shLZTR1. **d**, Loss of LZTR1 suppressed MAPK and CDK families of serine/threonine protein kinases as detected by Phospho-Threonine-Proline (P-Thr-Pro-101) antibodies. **e,** Validation of SKP2 downregulation in response to shLZTR1. These data exclude major roles for p21CIP1, p27KIP or p57 activation, known to participate in cell arrest, as the basis for LZTR1 growth suppression. Distinct shRNAs are numbered in **a-e**, as compared to control (scrambled) shRNA ('C'). Cell line identifiers are indicated above all plots in a and b and colored according to their origin: acral melanoma (blue), sun-exposed melanoma (red).

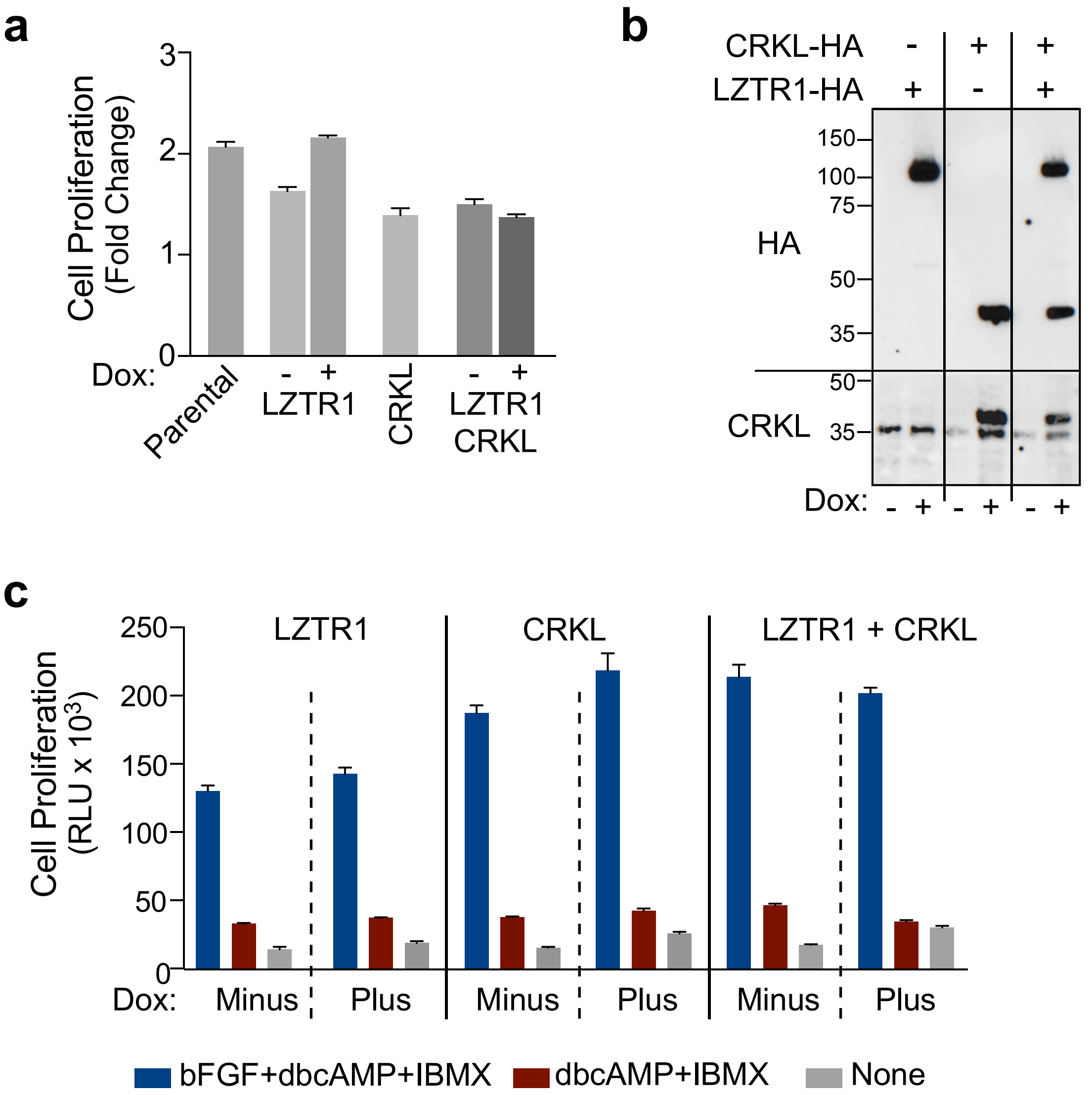


**Supplementary Figure 8: LZTR1 and CRKL, alone and in combination, failed to release human melanocytes from their dependency on growth factors. a,** Bar plots representing cell viability (CellTiter-Glo® Assay) in response to high expression of LZTR1-HA and PLX304-CRKL in normal human melanocytes (NBMEL C1220), as described in Fig. 6b. **b,** Western blot showing overexpression of LZTR1-HA and CRKL-HA, alone or in combination, in normal human melanocytes (NBMEL C454). **c,** Bar plots showing cell viability (CellTiter-Glo® Assay) performed after three days of incubation in the presence of the three growth factors, bFGF, dbcAMP, and IBMX (blue), the presence of just dbcAMP and IBMX (brown), or in the absence of all three factors (grey). Proliferation was reduced in all conditions relative to bFGF, dbcAMP, and IBMX, and in the absence of growth factors relative to dbcAMP and IBMX. Each measurement is the mean of triplicate (**a**) or duplicate (**c**) wells and error bars indicate SEM. Where indicated, medium was supplemented with 100 ng/ml doxycycline (Dox).
